# Chromatin Dynamics are Highly Subdiffusive Across Seven Orders of Magnitude

**DOI:** 10.1101/2025.05.10.653248

**Authors:** Matteo Mazzocca, Domenic N. Narducci, Simon Grosse-Holz, Jessica Matthias, Anders S. Hansen

## Abstract

Chromatin dynamics control the timescales of essential biological processes including DNA damage repair and activation of gene promoters by distal enhancers. Prior chromatin dynamics studies have reported widely varying degrees of subdiffusion, likely due to technical limitations. Here, we integrate MINFLUX—a recently developed single particle tracking method capable of achieving microsecond time resolution—with traditional tracking methodologies. We tracked chromatin dynamics across seven orders of magnitude in time in both human and mouse cells and found strongly subdiffusive dynamics (*α* ∼ 0.3). These dynamics are only mildly sensitive to perturbations of transcription, histone acetylation, and topoisomerase II, while loop extrusion has a constraining effect. Search times under these observed dynamics are extremely short for nearby loci (<100 nm), but almost impossibly long over larger distances (>1 µm). These findings have important implications for processes involving two locus contacts such as enhancer-promoter search and double-strand break repair.

## Introduction

Many essential processes within the nucleus require two pieces of chromosomal DNA to find each other. In mammalian gene regulation, distal enhancers must come into proximity with their cognate target genes to activate them, despite enhancers and promoters often being separated by tens to thousands of kilobases (*1*–*4*). Likewise, gene loci frequently locate near subnuclear structures including nuclear lamina, speckles, Polycomb bodies, and nucleoli, to regulate their activity or splicing (*5*–*7*). Similarly in DNA repair, the first step of both non-homologous end-joining and homologous recombination requires two loci to find each other to undergo repair (*8*). The timescale of these search processes depends critically on the dynamics of chromatin motion.

Chromatin motion is measured using live-imaging techniques, either at specific loci through DNA array reporters, or globally by fluorescently labeled histones (*9*–*11*). The observed dynamics are quantified by the mean squared displacement (MSD) as a function of time lag (*12*–*14*). In the field, MSD data are typically fitted to a power law, MSD(Δ*t*) ∝ Δ*t*^*α*^, for ease of analysis, modeling, and interpretation. The exponent *α* characterizes the nature of the observed motion: free diffusion (*α* = 1) describes a random walk with uncorrelated displacements, whereas subdiffusive behavior (*α* < 1) corresponds to constrained motion. The dominant constraint to chromatin motion is its polymeric nature, where adjacent nucleosomes are physically linked, leading to subdiffusion; for example *α* = 0.5 in the classical Rouse model of polymer dynamics (*15, 16*). This subdiffusive motion influences how quickly genomic elements locate one another: the lower the exponent *α*, the higher the tendency to explore the local surroundings, instead of the global environment. Accordingly, search times (the time required for two loci to find each other) become very short for proximal and excessively long for distal genomic elements. In mammalian nuclei, exponents ranging from 0.1 to 0.8 have been reported (Table S1). This broad spread in exponent values has substantial implications for search dynamics: for example, increasing the physical distance from 50 nm (∼10 kb) to 500 nm (∼500 kb; Eq. S52) increases the search time 300-fold if *α* = 0.8, but 10 billion-fold if *α* = 0.2 (Eq. S48). In terms of interactions between *cis*-regulatory elements, this could be the difference between distal interactions that are relatively frequent or impossibly rare over the course of a cell cycle, underscoring the need for understanding chromatin dynamics.

Precise determination of chromatin dynamics requires measurements both at very short timescales and over a large dynamic range. Crucially, the MSD at very short timescales strongly impacts search times at all scales. For example, a chromatin locus exhibiting a high *α* at short times may effectively miss targets below a specific size, having major implications for the kind of targets chromatin loci are capable of contacting. Additionally, while MSDs have conventionally been assumed to exhibit power law behaviors, accepting or refuting this in a robust manner requires a large dynamic range — at least 2 orders of magnitude (OOM) — in both time and space (*17*). Importantly, the smaller *α*, the larger the required time range. For example, if *α* = 0.4, as predicted by the fractal globule model (*18, 19*), > 5 OOM in time are required to get > 2 OOM in MSD. At an acquisition rate of 0.1 s, this would require 2.8 hours of imaging, which is highly challenging due to photobleaching. If instead *α* = 0.3, nearly 7 OOM are required, corresponding to 12 days of continuous imaging at the same acquisition rate (0.1 s). Due to the practical limitations of such imaging experiments, most prior studies do not have large dynamic ranges (Table S1). As a result, whether chromatin dynamics truly even exhibit power law behaviors remains unclear.

We overcame these challenges by integrating traditional tracking methods with MINFLUX, a recently developed single particle tracking technique capable of spanning over four OOM from hundreds of microseconds to seconds (*20, 21*). MINFLUX localizes particles through a successive triangulation approach which achieves extremely high resolution in both space and time, while minimizing photon flux (*20, 21*). While MINFLUX has been successfully used to track freely diffusing and processive proteins (*20, 22*–*27*), it has not yet been validated for tracking chromatin in living cells. Here, for the first time, we applied MINFLUX tracking to study chromatin dynamics in live mammalian cells and integrated MINFLUX with conventional single-molecule histone tracking (*28*) and super-resolution live-cell imaging of DNA loci (*29*). This multi-modal approach allows us to study chromatin dynamics at sub-millisecond timescales and over seven orders of magnitude in time.

## Results

### MINFLUX for measuring chromatin dynamics

To measure chromatin dynamics across seven orders of magnitude (OOM), we integrated three different live-cell fluorescence microscopy techniques that allow tracking chromatin dynamics (Fig. **1A**). To overcome previous limitations at very short timescales, we applied MINFLUX (*20*) to track chromatin dynamics across four OOM (200 µs – 10 s). For intermediate timescales (100 ms – 200 s) we employed camera-based single-particle tracking (SPT) (*28*), and for longer timescales we used Super-Resolution Live-Cell Imaging (SRLCI) data (20 s to hours) (*29*). These methods cover partially overlapping timescales, thus allowing for cross-method comparisons.

**Figure 1:**
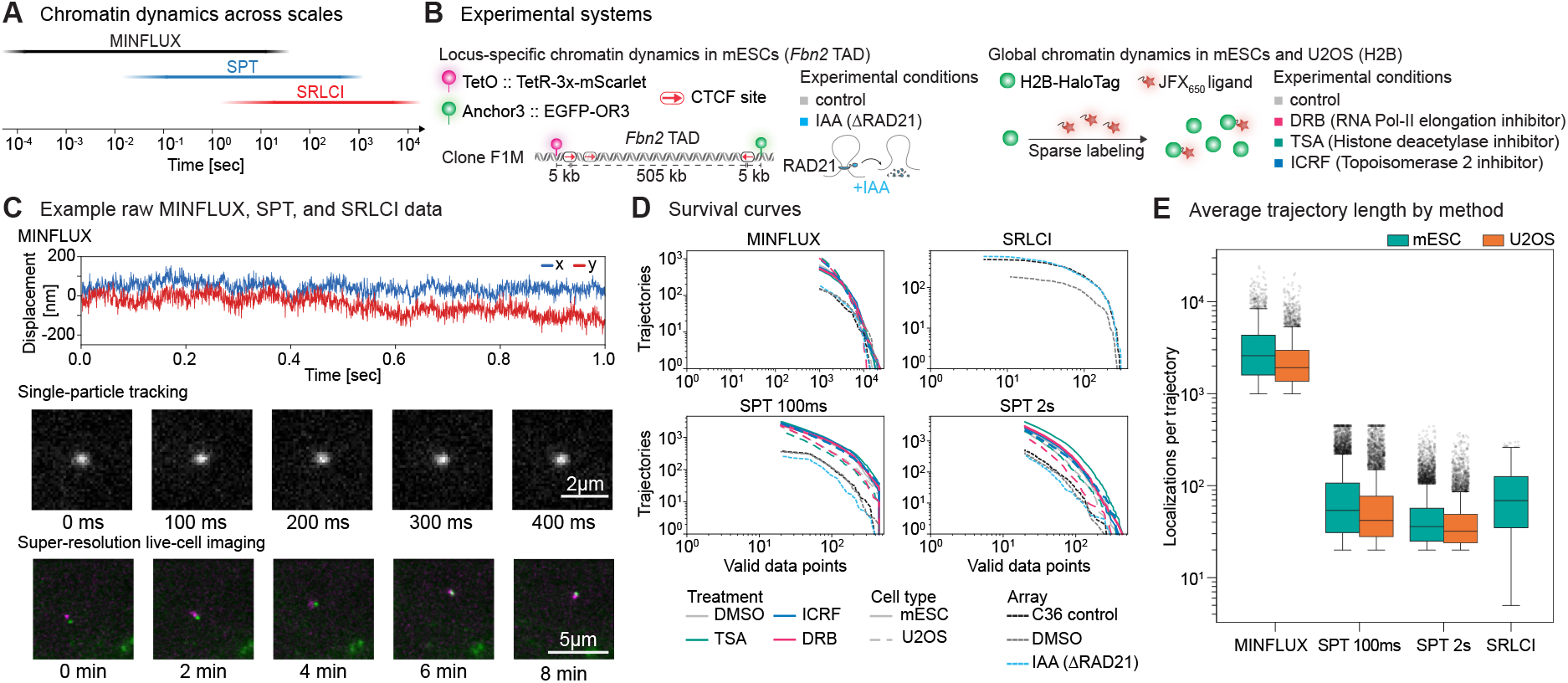
Multimodal live-cell microscopy approach to track locus-specific and global chromatin dynamics across seven orders of magnitude. (A) Three complementary approaches to track chromatin motion across timescales in living cells: MINFLUX (black line) for short timescales (200μs–ls), single-particle tracking (SPT, red line) for intermediate timescales (100 ms–200 s), and super-resolution live-cell Imaging (SRLCI) for long timescales (20 s–hours). (B) Experimental system to study locus-specific and global chromatin dynamics and associated conditions. On the left, a diagram of the mESC line showing the arrangement of fluorescent reporter arrays at the Fbn2 TAD for locus tracking. On the right, a diagram of the labeling strategy for the H2B-Halo in mESCs and human U2OS cells. (C) Raw data from different tracking approaches. 2D MINFLUX tracks particles as (*x,y*) coordinates over time (top), while camera-based methods localize them a posteriori in recorded movies: H2B-Halo (SPT, middle), DNA arrays flanking the Fbn2 TAD (SRLCI, bottom). (D) Survival curves of chromatin trajectories across microscopy approaches and treatment conditions. Due to higher photobleaching in SPT, we used two acquisition settings (100 ms or 2 s frame interval) to cover three orders of magnitude. (E) Number of localizations per trajectory for each approach. Outliers are indicated by transparent black dots.

We designed an experimental scheme to enable three comparisons: 1) locus-specific versus global chromatin dynamics; 2) chromatin dynamics between mouse and human cells; 3) response to perturbations of key chromosomal processes (Fig. **1B**). To study chromatin dynamics at a specific locus, we used previously validated mouse embryonic stem cell lines (mESC) (*29*). These lines (C36 and F1M) contain fluorescent reporter arrays at the CTCF boundaries of the 505 kb *Fbn2* TAD. Additionally, the cohesin subunit RAD21 is mAID-tagged in F1M, allowing for acute degradation upon treatment with indole-3-acetic acid (IAA). To measure chromatin dynamics throughout the genome in two different species, we used mESC and human U2OS cell lines that stably express Halo-tagged histone H2B, which can be labeled with fluorescent ligands (*30, 31*). To perturb chromatin motion we used drugs known to inhibit chromatin-related molecular processes, specifically: transcriptional elongation by RNA-Pol II, histone deacetylation, and topological relaxation by Topoisomerase II (*28, 32*–*34*).

Unlike SPT and SRLCI, which perform localization on images, MINFLUX directly tracks single particles without imaging and outputs a set of coordinates over time (Fig. **1C**). MINFLUX requires 20-100 times fewer photons than camera-based localization to achieve the same precision, thus increasing the information content which can be extracted before photobleaching (*20*). With MINFLUX, therefore, we collect on average 3.5 times more localizations per trajectory than SRLCI and 8-20 times more than SPT (Figs. **1D** and **1E**).

Because it has so far never been used for chromatin tracking, we first characterized MINFLUX tracking (Abberior implementation; Supplementary Information 2.3). MINFLUX estimates particle positions using a donut-shaped beam with zero intensity at the center. Each iteration involves six or seven probings at different donut positions, forming a hexagonal pattern of diameter L (Fig. **2A**). The recorded photon numbers at each probing position, together with the known non-uniform intensity of the beam, enable the precise triangulation of the particle position (*21*). After determining the molecular coordinates, the donut moves to this position, starting the next probing cycle and allowing it to track H2B-Halo or the *Fbn2* TetO array at 200 µs-resolution (Fig. **2B** and Fig. S1).

**Figure 2:**
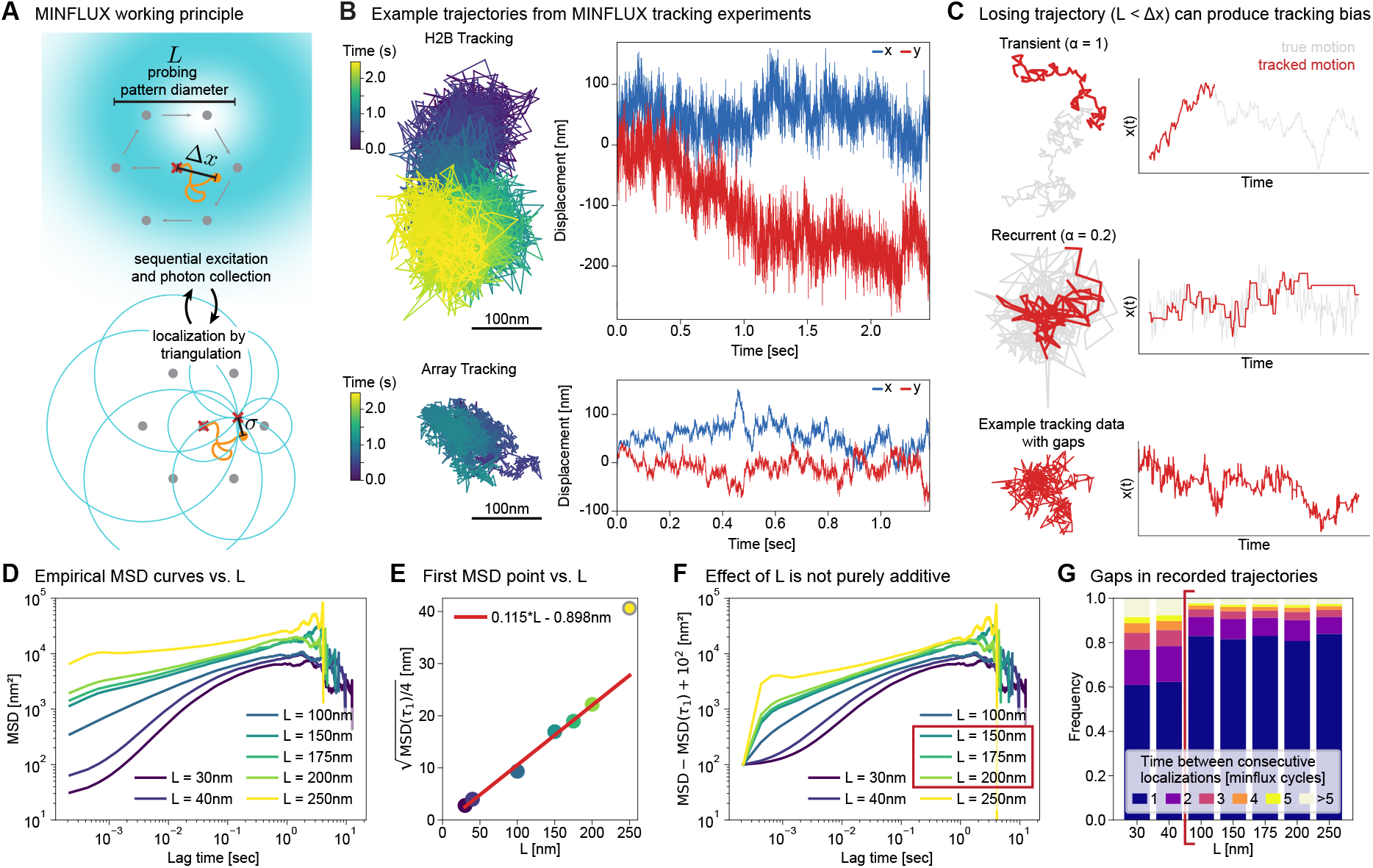
MINFLUX working principle and effect of probing pattern diameter on tracking output. (A) MINFLUX (Abberior implementation) excites the particle with a donut-shaped beam with zero intensity at the center. A single localization requires a probing sequence (iteration) with a hexagonal pattern of diameter *L*. To track the particle over time, the donut moves to the estimated particle position, initiating the next iteration. (B) MINFLUX 2D tracking of H2B-Halo (top) and *Fbn2* locus (TetO array, bottom). On the left, trajectories depict particle motion in space, with color indicating tracking time. On the right, corresponding spatial coordinates over time (*x, y* in blue, red, respectively). (C) Simulations of true particle motion (gray) and biased tracking (red), where only particle locations within a region of size *L* around the last known localization are tracked, for normal diffusion (*α* = 1, top) and recurrent motion (*α* = 0.2, middle). The latter shows pronounced gaps in tracking and an associated bias against large displacements. Bottom: real MINFLUX trajectory for comparison. (D) MSD curves from MINFLUX tracking of H2B-Halo in U2OS cells, using different probing pattern diameters *L*. (E) First MSD point of H2B-Halo for different *L*, showing linear scaling (expected for localization error). (F) MSD curves after subtracting the first MSD point (and adding a constant offset for visualization), indicating inconsistency across *L* is not solely due to additive terms such as localization error. (G) Distribution of temporal gaps in trajectories, dependent on *L*.

The probing pattern diameter *L* is a central MINFLUX acquisition parameter that must be optimized, specifically for tracking applications. There is a tradeoff between decreasing *L*, which reduces localization error (*20*), and increasing *L*, which reduces tracking bias. Tracking bias arises from gaps in trajectories due to intermittent particle loss when the particle’s step size Δ*x* between localizations is comparable to *L*; akin to using too small a search range in camera-based tracking algorithms. To evaluate this tracking bias, we simulated MINFLUX tracking for free diffusion (*α* = 1) and strongly subdiffusive fractional Brownian motion (fBm, *α* = 0.2; Fig. **2C**) (*12*). Freely diffusing particles exhibit few tracking gaps: once lost, the particle does not return and remains undetectable. Fractional Brownian motion, in contrast, becomes recurrent for *α* < 2*/d* = 0.66 (SI; *d* = 3 being space dimension), meaning that if the particle takes a large step and escapes the excitation region, it will return and continue being tracked. The resulting gaps constitute a bias against recording large displacements (Fig. **2C**). Choosing *L* too small is thus expected to strongly decrease the observed particle mobility at short times, if particle motion is recurrent (small *α*), thereby introducing bias.

To optimize *L* for chromatin tracking, we tracked chromatin motion in U2OS cells using MINFLUX for *L* between 30 and 250 nm (Fig. **2D**). As expected, the observed MSD over short time lags is strongly affected by the choice of *L*, due to localization error (increasing with *L*) and bias (decreasing with *L*). An upper bound on localization error can be obtained from the first MSD point and was found to scale linearly with *L* (with *L* = 250 nm as an outlier; Fig. **2E**). We next assessed whether the difference between the MSD curves could be explained by localization error alone, by subtracting the first MSD point (Fig. **2F**). While for *L* = 150, 175, and 200 nm, MSD curves approximately collapsed, the curves for *L* = 30, 40, 100, and 250 nm remain qualitatively different, presumably due to bias for small *L* and instrument limitations for 250 nm. We next quantified temporal gaps in the trajectories and found a vast increase in gaps for *L* = 30, 40 nm (Fig. **2G**). The gap distribution stabilized around *L* ∼ 100 nm. Combining our insights from MSD curves and gap distributions, we decided to use *L* = 150 nm for subsequent experiments. This choice leads to a localization precision of approximately 15 nm (Table S2) while minimizing bias due to intermittent particle loss, thus allowing us to track chromatin motion with MINFLUX for the first time (Fig. S2–5).

### Chromatin dynamics across many OOM

To integrate observations of chromatin motion by MINFLUX, SPT, and SRLCI (Fig. S6–15), we need to account for technical differences between these techniques, including localization error, motion blur bias, cell motion, spatial dimension (2D vs. 3D tracking), and number of loci tracked. To correct for cell motion, we calculated MSDs in SPT and SRLCI as 0.5 times 2-point MSD (SI). The remaining method-dependent error terms can cause pronounced mismatches between observed MSDs, even for data that are synthetically generated from a single fBm process (Fig. **3A**), making it visually difficult to judge whether MSDs measured by different techniques are consistent with each other. We therefore plot three sets of curves when discussing MSDs: the observed MSD for each technique (blue); the proposed underlying true MSD (orange; e.g. a fit); and how this true MSD would be distorted by each tracking technique (red; Fig. **3A**, right panel). For fitting analytical MSD curves to data, we extended a previously developed Bayesian approach (*29, 35*) (SI), allowing us to jointly fit data from different tracking approaches while taking into account the technical distortions associated with each. This approach accurately recovered the true MSD from the synthetically distorted data (Fig. **3A**).

**Figure 3:**
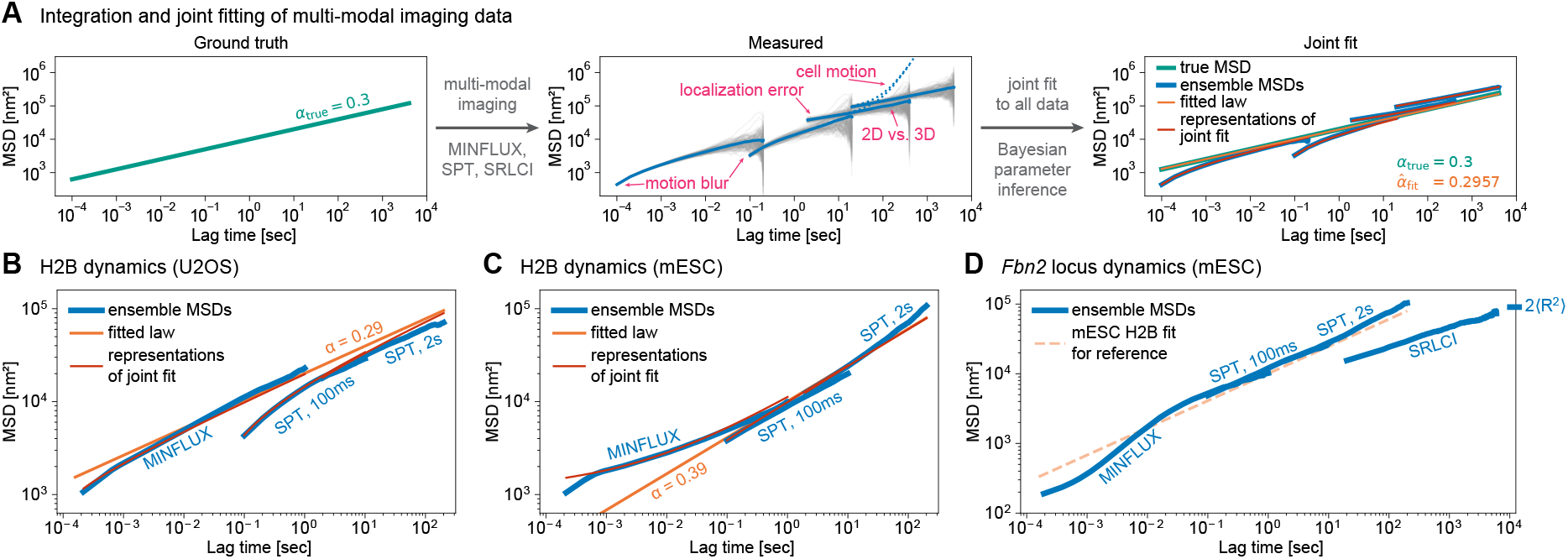
Integrating MINFLUX, SPT, and SRLCI to study chromatin dynamics. (A) Schematic demonstrating how technical sources can distort data from a simple power law MSD. Left: ground truth used to generate trajectories. Middle: synthetic data following the ground truth, plus indicated error sources. Right: jointly fitting all data accurately recovers the ground truth MSD. Red lines indicate expected MSD given ground truth and fitted/known error terms. (B, C) Ensemble MSDs (blue) with the fitted power law MSD (orange) and corresponding dataset-specific fit lines (red). (B) U2OS cells, (C) mESCs. (D) Ensemble MSDs (blue) for the *Fbn2* locus in mESC clone C36, with joint fit from mESC H2B-Halo tracking (C) shown for reference (orange dashed). The line at the axes border indicates limiting MSD value for SRLCI, given by the average separation between the two loci.

We next applied this method to our histone tracking data in U2OS cells (Fig. **3B**) and mESCs (Fig. **3C**). The U2OS data is well represented by a single power law MSD with an exponent of 0.29, the observed “downward curving” MSDs being consistent with the effect of motion blur for both MINFLUX and SPT (Fig. **3B**). Fitting each of the data sets (MINFLUX, SPT at 100ms, SPT at 2s) separately or in various combinations yielded consistent results (Fig. S16– 19). We similarly attempted to fit a single power law to the mESC data (Fig. **3C**). In this case, however, the fit converged to a significantly higher localization error in the MINFLUX data set (Table S2), to account for the “upward curve” at short times. Fitting each data set individually furthermore gave inconsistent results, with the best-fit exponent increasing from 0.32 for MINFLUX, to 0.42 for SPT (100 ms), to 0.60 for SPT (2 s) (Table S2 and Fig. S15). We therefore conclude that chromatin dynamics in human U2OS and mouse ES cells are fundamentally different: chromatin dynamics in U2OS cells exhibit power law scaling across the full dynamic range with exponent *α* ≈ 0.29, whereas chromatin dynamics in mESCs are not faithfully represented by a single power law MSD, invalidating the use of a single “exponent” to characterize mESC dynamics.

We then proceeded to study chromatin dynamics at the *Fbn2* locus in mESCs, extending the existing long-timescale SRLCI data (*29*) to intermediate and short time scales using SPT and MINFLUX, respectively (Fig. **3D**). Relative to mESC histones, we noted a marked decrease of mobility at short timescales, which we attribute to tracking a finite size array (10 kb) instead of single histones. The localization of this array (center of emission) is expected to diffuse within the fluorescent mass of the array at short times (MINFLUX), leading to the observed drop in MSD. At intermediate time scales (SPT) the observed MSD is remarkably consistent with the mESC histone tracking, while the long-time scale (SRLCI) data deviates markedly from that trend. We attribute the latter deviation to the fact that the two loci tracked in SRLCI are genomically close to each other (SI 3.3) (*29*).

### Perturbations of chromatin dynamics

Lastly, we performed H2B and *Fbn2* locus tracking under treatment conditions to understand how transcription, histone acetylation, topological effects, and loop extrusion affect chromatin dynamics (Fig. **4A**). We first treated both U2OS cells and mESCs with either DRB, an RNA Polymerase II inhibitor (*36*), TSA, a histone deacetylase inhibitor (*37*), or ICRF, a topoisomerase II inhibitor that increases genomic supercoiling (*38*). In U2OS cells, we find that inhibiting transcription with DRB increases chromatin mobility with no meaningful change in exponent, consistent with prior work (*28, 32, 39, 40*) while inhibiting topoisomerase II causes a similarly sized decrease in chromatin mobility (Fig. **4B**). Increased histone acetylation upon TSA treatment slightly increased chromatin mobility in U2OS cells. This result was largely consistent (albeit with smaller effect size) in mESCs, for which DRB treatment caused a minor increase in mobility and ICRF a slight decrease (Fig. **4C**). However, compared to U2OS, in mESCs, TSA treatment modestly increased chromatin mobility, consistent with prior work in other cell types (*28, 41*). Finally, we examined loop extrusion by cohesin by tracking *Fbn2* locus dynamics in the presence or absence of RAD21 (*29*) (Fig. **4D**). At long time scales (SRLCI), degradation of cohesin increases chromatin mobility and steady-state distance between the two tracked loci (*28, 29, 42*). At intermediate time scales (SPT), a slight increase in mobility relative to the control remains. At the shortest time scales (MINFLUX) we do not observe a difference between degron and control, since the dynamics are dominated by the internal fluctuations of the 10-kb array. Notably, we did not observe major qualitative changes in MSD (e.g. in exponent for power laws) upon any perturbation except for ΔRAD21. This suggests that loop-extrusion is a primary regulator of chromatin motion on time scales of minutes to hours.

**Figure 4:**
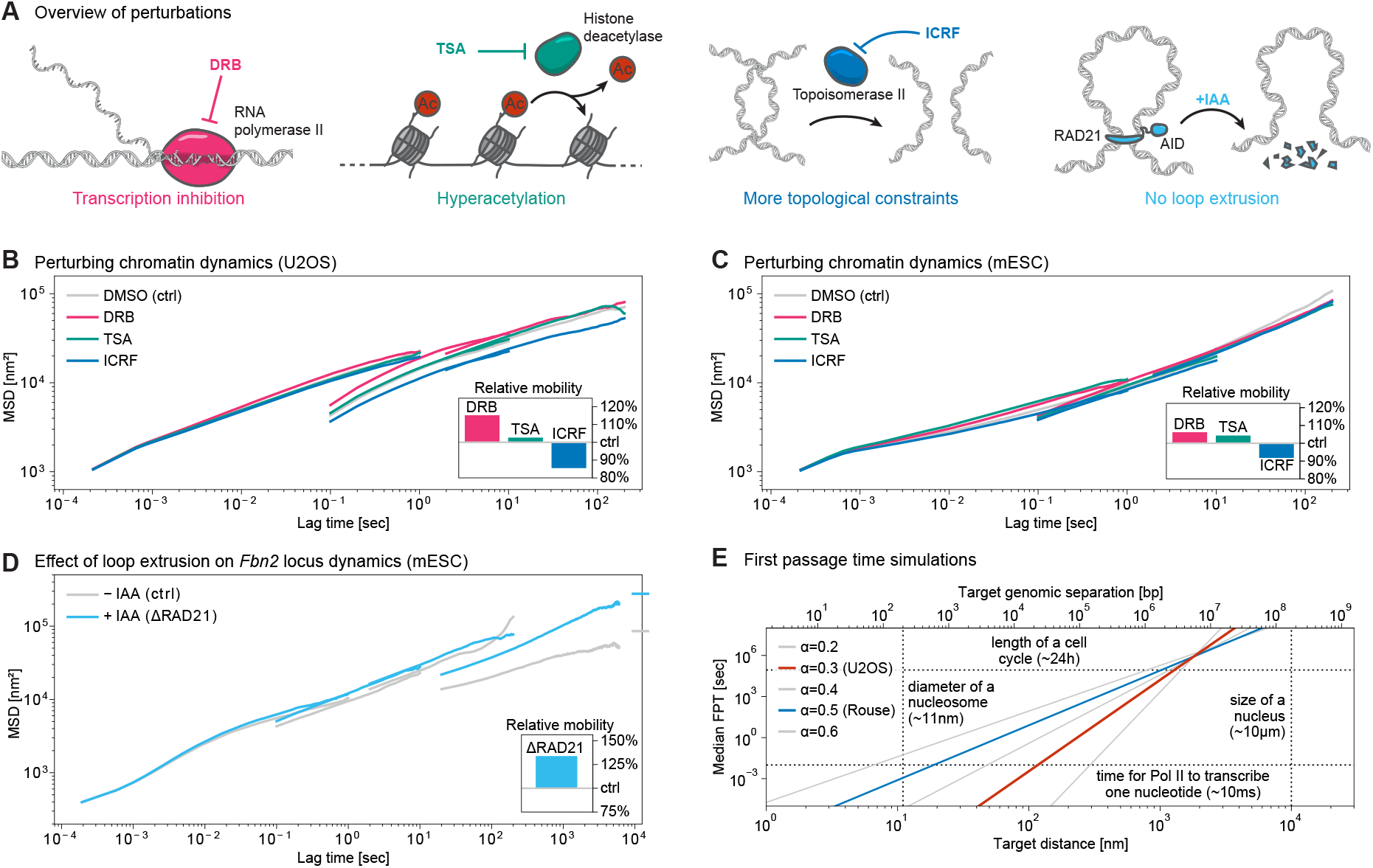
Chromatin dynamics are largely invariant to perturbations of transcription, histone acetylation, and topoisomerase II, but constrained by loop extrusion. (A) Schematic of the drug treatments used in this study and their effect. From left to right: DRB inhibits RNA polymerase II, TSA inhibits histone deacetylase, ICRF inhibits topoisomerase II, and RAD21 depletion abolishes loop extrusion. (B) Ensemble MSDs of H2B-Halo in U2OS cells for each of the treatment conditions (gray: control treatment with DMSO; magenta: DRB (3h); teal: TSA (4h); blue: ICRF (2h)). The average mobility relative to the control condition is inset in the bottom right. (C) Ensemble MSDs of H2B-Halo as in (B) but in mESCs. (D) Ensemble MSDs from the locus tracking experiments as in (B) (gray: untreated condition; cyan: RAD21 depletion (2h + 4h)). Lines at the axes border indicate limiting MSD value for SRLCI data, calculated as 2 ⟨*R*^2^ ⟩, *R* being the absolute distance between the loci. (E) First passage times as a function of target distance for a point-like target. MSD exponents of 0.2, 0.4, and 0.6 are shown in light gray, while 0.3 (resembling U2OS), is emphasized in red and 0.5 (Rouse dynamics) is emphasized in blue. Relevant length and time scales are shown as dashed lines.

## Discussion

Integrating MINFLUX with SPT and SRLCI allowed us to study chromatin dynamics over an unprecedented seven OOM. Strikingly, over seven OOM, from 200 µs to hours, the MSD only increases about two OOM. Converting from MSD to linear displacements, this means that a DNA locus only moves ∼ 300 nm in 2D and ∼ 500 nm in 3D over these timescales. While histone dynamics in U2OS cells are well represented by a single power law with an exponent 0.29, chromatin motion in mESCs seems to be qualitatively different on different time scales, with the MSD becoming steeper at longer times. This absence of a universal power law scaling in mESCs might explain the diversity of exponent values reported in the literature (Table S1): the estimated “exponent” will depend on the timescale under study. This change in chromatin motion over timescales opens the questions of what are the underlying driving mechanisms of such change, and the resulting implications on the search times. Regardless of the precise analytical form (power law or not), MSDs in both U2OS and mESC are very shallow—and indeed shallower than most previous reports (Table S1)—indicating strong subdiffusion.

What do such strongly subdiffusive dynamics imply for two DNA loci trying to find each other in the nucleus? Subdiffusion can be thought of as a trade-off between global versus local exploration. Highly subdiffusive motion means that the local environment is very quickly and repeatedly searched, at the cost of undersampling globally. To give quantitative intuition for this behavior, we simulated fractional Brownian motion (fBm) with MSD exponents *α* ranging from 0.2 to 0.6 and calculated median first passage times (FPT) to a point-like target at a distance *X*; this is a self-similar problem and governed by the scaling relation FPT ∝ *X*^2/*α*^ (Fig. **4E**, SI 5.1, and Fig. S20). For small MSD exponent *α*, this FPT scaling becomes very steep: for the reference simulation resembling our U2OS data (*α* = 0.3), a median FPT of 10 seconds corresponds to an initial separation of 330 nm between the two loci. Requiring faster (100 ms) or allowing for slower (1000 s) target search shifts this required initial separation only to 170 or 660 nm, respectively. Thus, the limit for successful search in the nucleus—over biologically relevant time scales—is an initial separation of the two loci of hundreds of nanometers. Strongly subdiffusive chromatin dynamics makes encounters with the immediate surroundings (<100 nm) extremely frequent (<4 ms) and very distal interactions (>1.3 µm) nearly impossible (>25 h; Fig. **4E**).

These findings have major implications for any process requiring two pieces of DNA to find each other, including enhancer-promoter (E-P) interactions and DNA repair. For double-stranded DNA breaks, highly subdiffusive dynamics of the loose DNA double-strand break ends will keep them in proximity until repair machinery arrives (*8*). For distal gene regulation, a major unsolved question is how close in space an enhancer needs to be to activate its target promoter (*2*). Classically, E-P loops were thought to require physical contact (<10 nm), though recent papers have proposed action-at-a-distance models involving gradients or condensates (>200 nm) (*2, 43*–*46*). A direct consequence of highly subdiffusive chromatin dynamics is that within ∼200 nm, diffusive search is so efficient (∼360 ms) that frequent physical contact is inevitable; the distinction between “contact” and “action-at-a-distance” (up to ∼200 nm) then ceases to be meaningful. Thus, our findings may reconcile the seeming contradiction between classical “activation-by-recruitment” models involving close contact (*47, 48*) and more recent measurements of larger E-P distances during transcriptional activation (*2, 43*–*46, 49*).

With hundreds of genes and thousands of enhancers per µm^3^ in the nucleus (*2*), subdiffusive chromatin dynamics imply frequent, all-to-all encounters for close-by elements: within a radius of 300 nm, a given promoter can find tens of genes and hundreds of enhancers within minutes. Indeed, multi-way, all-to-all E–P and P–P interactions spanning hundreds of kb, termed microcompartments, have been observed in gene-rich chromatin regions (*50, 51*). Given this promiscuity, selectivity in functional, short-range E– P interactions cannot be achieved through the search process alone, but likely require other mechanisms such as biochemical E-P selectivity (*2*). A tenfold increased E–P separation, on the other hand—for example 1 Mb instead of 100 kb—increases search time ∼5 million fold (to ∼5 h) if *α* ≈ 0.3, making robustly successful target search virtually impossible by baseline chromatin motion alone. Productive interactions between distal E–P pairs thus require facilitation by processes such as loop extrusion (*52*). For example the oncogene *MYC* relies on CTCF-anchored loop extrusion to interact with enhancers up to 1.8 Mb away (*53, 54*).

In summary, we integrated MINFLUX, SPT, and SRLCI data to study chromatin dynamics over seven OOM in time. We applied and validated MINFLUX for the use of chromatin tracking in live cells for the first time. While dynamics in U2OS cells are well represented by a single power law with exponent 0.29, our data suggest a continually steepening MSD for mESCs, potentially explaining inconsistent “exponent” values in the literature. This behavior was mildly affected by perturbations of chromatin-related processes. Our determination of chromatin dynamics with unprecedented dynamic range—arguably covering most if not all time scales relevant to cycling cells—provides a foundational building block in our spatio-temporal understanding of the cell nucleus.

## Acknowledgements

We thank Luca Giorgetti, Davide Michieletto, Christoph Zechner, Edward Banigan, Jamie Drayton, Michele Gabriele, Clarice Hong, Miles Huseyin, Christos Katsifis, Sumin Kim, Masahiro Nagano, Henrik Pinholt, Varshini Ramanathan, Jack Toppen, and Harvey Yang for generously providing feedback on the manuscript, and we thank the Hansen, Mirny, Zechner, and Brugués labs for helpful discussions. We thank Luke Lavis for providing Janelia Fluor dyes. We thank Karsten Bahlmann for coordination of the project, and we thank Murali Palangat and Tatiana Karpova for hosting the MINFLUX experiments at the NIH.

## Author contributions

All authors contributed to study design and conceptualization. Experiments were performed by MM, JM, DN (MINFLUX) and MM (SPT). MM tracked SPT data. Data processing, refinement, and analysis was performed by SGH. MINFLUX and fBm simulations were run by DN, SGH contributed to analysis. ASH, MM, DN, SGH drafted the manuscript. All authors edited and approved the manuscript.

## Data and material availability

Cell lines, plasmids, and other materials are available upon request. The code and software used in this study are freely available on GitHub (https://github.com/ahansenlab/chromatin_dynamics).

The raw trajectory data are available through Zenodo (DOI:10.5281/zenodo.15369544).

