## Supplementary Information for "Chromatin Dynamics are Highly Subdiffusive Across Seven Orders of Magnitude"

May 10, 2025

#### Materials and Methods

**1 Cell culture, labeling, and treatment conditions** **3**

**6 Supplementary Figures** **16**

**7 Supplementary Tables** **36**

**8 Supplementary Movies** **42**

**9 References** **43**

### 1 Cell culture, labeling, and treatment conditions

#### 1.1 Cell culture

To study histone dynamics we used human U2OS and mouse embryonic stem cells (mESCs, JM8.N4 (*1*)) stably expressing H2B-Halo (*2*). For locus tracking, we used mESCs (JM8.N4) carrying the TetO and Anchor3 arrays at the *Fbn2* locus, stably expressing fluorescent proteins that bind these arrays (TetR-3x-mScarlet and EGFP-OR3, respectively) and endogenously expressing CTCF-Halo (cell lines C36 and F1M) (*3*). In addition, cell line F1M expresses RAD21-tagBFP-mAID-V5 and OsTIR1 allowing rapid RAD21-depletion follwoing treatment with indole-3-acetic acid (IAA) (*3*). All mESC lines employed in this study were cultured on plates pre-coated with 0.1% sterile gelatin solution (Sigma-Aldrich, G1890) under feeder-free conditions. MESCs were grown 37℃ and 5% $CO_2$, in a medium consisting of KnockOut™ DMEM (ThermoFisher, 10829-018) supplemented with 15% FBS (Avantor Seradigm, 89510-186, Lot 190B20) 1000 U/mL LIF homemade) (*2*), 1 mM MEM Non-Essential Amino Acid Solution (ThermoFisher, 11140050), 2 mM GlutaMAX™ (ThermoFisher, 35050061), 100 µg/mL Penicillin-Streptomycin (ThermoFisher, 15140122), 0.1 mM 2-mercaptoethanol (ThermoFisher, 31350010), 10 µM MEK inhibitor PD0325901 (Tocris, 4192), and 3 µM GSK-3 inhibitor CHIR99021, (Sigma-Aldrich, SML1046).

Human U2OS cells were cultured at 37℃ and 5% $CO_2$ in DMEM (ThermoFisher, 31053-028) supplemented with 10% FBS (Avantor Seradigm 89510-186, Lot 190B20), 2 mM GlutaMAX™ (ThermoFisher, 35050061), 100 µg/mL Penicillin-Streptomycin (ThermoFisher, 15140122), and 100 nM Sodium Pyruvate (ThermoFisher, 11360070).

#### 1.2 Cell seeding and labeling for live-imaging experiments

For SPT, mESCs and U2OS cells expressing H2B-Halo, and the mES cell lines C36 and F1M for *Fbn2* locus tracking, were seeded the day before imaging on 35 mm glass-bottom dishes (MatTek Life Sciences, P35G-1.5-14C). For MINFLUX, the same cell lines were seeded on 8-well Nunc™ Lab-Tek™ glass-bottom chambers (ThermoFisher, 155409PK). Before seeding mESCs, imaging dishes or chambers were coated with MatriGel (Corning, 354277) following the manufacturer's instructions. mESCs were grown overnight in KnockOut™ DMEM supplemented as described in the cell culture section. On the imaging day, KnockOut™ DMEM was replaced with phenol red-free DMEM (ThermoFisher, 31053-028), keeping all other medium components unchanged. Prior to imaging, H2B-Halo mESCs were labeled by adding two fluorescent Halo-ligands to the medium: one at a relatively high concentration to obtain a reference image of the nucleus ($JFX_{554}$ at 100 pM for SPT or 50 pM for MINFLUX) and the other at a single-molecule density for tracking H2B dynamics (2.5 pM $JFX_{650}$). For SPT of the *Fbn2* TetO array, C36 and F1M cells were incubated with the Halo-dye $JFX_{650}$ (2.5 pM) to label CTCF-Halo. This allowed us to track both *Fbn2* (through the constitutive expression of TetR-3x-mScarlet3) and CTCF-Halo, the latter of which was used for cell motion correction (see Section 3.2.2). mESCs labeling was performed by incubating cells with Halo-ligands at 37℃ for 30 minutes, followed by three washes in culture medium, a 30-minute incubation at 37℃ for further washout, and a final replacement with fresh medium.

H2B-Halo U2OS cells were grown in phenol red-free DMEM as described in the cell culture section. Before imaging, labeling and washout were performed as described for H2B-Halo mESCs using two fluorescent Halo-ligands but at different concentrations: $JFX_{554}$ to map the cell nucleus (10 pM for SPT or 50 pM for MINFLUX) and $JFX_{650}$ for tracking H2B dynamics (0.5 pM for SPT or 2.5 pM for MINFLUX).

#### 1.3 Chemical treatments

Treatments to perturb chromatin dynamics were performed before live-imaging experiments and maintained during imaging, for a total duration of up to 2 hours from the start of treatment. mESCs and U2OS cells expressing H2B-Halo were treated prior to imaging for 3 hours with 100 µM DRB (Cayman Chemical, 10010302) to inhibit transcription (*4*), for 2 hours with 5 µM ICRF-197 (Enzo Life Sciences, BML-GR332) to inhibit DNA topoisomerase II activity (*5*)(*6*), and for 4 hours with 150 nM, 500 nM, 1.5 µM TSA (Sigma-Aldrich, T8552) to inhibit histone deacetylase (*7*). The F1M cell line used for *Fbn2* locus tracking and expressing RAD21-tagBFP-mAID-V5, was treated prior to imaging for 3 hours with 500 µM IAA (Santa Cruz Biotechnology, sc-254494) to induce RAD21 degradation. All chemicals were added to the culture media from 1000x stock solutions in DMSO. Control samples for mESCs and U2OS cells expressing H2B-Halo, and the F1M cell line were treated with 0.1% DMSO for up to 4 hours before imaging, matching the longest treatment condition (TSA).

#### 2 Microscopy experiments and analysis

##### 2.1 Single-particle tracking (SPT) to track Histone H2B-Halo and *Fbn2* locus dynamics

SPT experiments were performed on a custom-built (*8*) Nikon Ti2-E inverted microscope with a 100x, 1.49 NA oil immersion apochromatic objective and two Prime 95B sCMOS cameras (Teledyne Photometrics, 110 nm pixel size), equipped with 593/40 nm and 698/70 nm bandpass filters, respectively. HILO illumination was achieved by controlling the excitation laser angle using an iLas 2 motorized dual galvo system (Gataca Systems). The inclination was set at 57° and the illumination in HILO ellipse mode, where the lasers spin in an ellipse (6.67 ms per spin), providing uniform illumination across the field of view and enabling simultaneous imaging of multiple cells, unlike regular HILO where the laser beam crosses only one cell at a time (*9*). The sample conditions were kept at 37℃ with 5% $CO_2$ and controlled humidity using an Okolab stage-top chamber. To track H2B-Halo, samples were excited with a 1W 561 nm Coherent Genesis laser at 5% power for the first and last frame to obtain a reference image of the nucleus, and with a 1W 640 nm Coherent Genesis laser at 4% (U2OS) or 6% (mESCs) for SPT movies. To track both the *Fbn2* TetO array and CTCF-Halo (used for for cell motion correction; Section 3.2.2) in C36 and F1M cells, we performed dual-color SPT using the 1W 561 nm Coherent Genesis laser at 5% to excite TetR-3x-mScarlet3, and the 1W 640 nm laser at 10% for CTCF-Halo. Laser power was modulated by an acousto-optic tunable filter (AAOpto-Electronic, AOTFnC-400.650-TN) triggered by the camera exposure TTL signal. The imaging ROI was set to 512 × 512 pixels, allowing multiple cells to be imaged simultaneously. To capture the full range of H2B and *Fbn2* dynamics covered by SPT in this study (from 100 ms to $\tilde{1}$0 minutes), we established two SPT acquisition regimes to sample both fast and slow dynamics while minimizing signal loss from photobleaching. We applied a fast regime with a 100 ms lag time between adjacent frames (camera exposure: 100 ms/frame 10 fps) acquiring 450 frames, and a slow regime with a 2 s lag time (camera exposure: 2 s/frame, 0.5 fps) acquiring 450 frames (H2B-Halo) or 300 (*Fbn2*/CTCF-Halo). In both regimes, the excitation laser was pulsed for 86.71 ms per frame (corresponding to 13 cycles of the HILO ellipse mode), blurring out non-chromatin-bound molecules and ensuring a comparable photobleaching rate in both regimes.

##### 2.2 Image processing of SPT movies to obtain H2B-Halo and *Fbn2* locus trajectories

Since the large field of view (512 × 512 pixels) often included multiple cells, we manually drew an ROI around each nucleus to ensure that H2B or *Fbn2*/CTCF-Halo particles were analyzed independently for each cell. As described in the previous section, H2B-Halo SPT acquisitions consisted of two reference images of nuclei (first and last frames in the 561 nm channel) and the SPT movie itself in between (640 nm channel). Since cells could move during acquisitions, the two reference images were compared to draw ROIs that overlapped in both the first and last frame while avoiding neighboring nuclei. Similarly, for *Fbn2*/CTCF-Halo, the first and last frames of the movie were compared using the TetR-3x-mScarlet3 signal (561 nm channel), which provides a good reference image of nuclei. Additionally, *Fbn2* movies were manually inspected to identify potential replicated sister chromosomes, appearing as two overlapping fluorescent dots. Cells with replicated chromosomes were discarded.

SPT movies were subsequently analyzed using the TrackMate plugin in ImageJ, which detects individual spots in each frame and links corresponding spots across frames into continuous tracks (*10*). Single particles (corresponding to H2B-Halo molecules, *Fbn2* TetO array, or CTCF-Halo molecules) were identified by applying a Laplacian of Gaussian (LoG) filter to each frame, with a maximum particle diameter of 0.8 µm and a minimum quality score of 3. This quality score reflects spot brightness and how closely the spot size matches the specified diameter. Detected spots were then linked into trajectories using the LAP algorithm (*11*), which requires specifying a maximum allowed displacement between frames for each detected particle. The maximal frame-to-frame displacement for single particles was set to 300 nm and 450 nm for 100 ms and 2 s lag time movies, respectively. These thresholds were determined by measuring the frame-to-frame displacement of detected particles at frame 0 and comparing them with those at frame 1 in a preliminary subset of movies. The displacement distributions of "all-vs-all detections" were then plotted and used to inform these thresholds. Finally, TrackMate generated tables containing the $x$, $y$ coordinates and time points for each track, along with a tracking model that could be reloaded into TrackMate to visualize ROIs and particle tracks overlaid on the movies.

A few mESCs movies containing dead cells or cells imaged too close to the coverslip were manually excluded from analysis: movie 000 in DMSO (control), 100 ms, replicate 3; movie 000 in ICRF, 100 ms, replicate 3; and movies 000, 001, 002, and 003 in DMSO (control), 2 s, replicate 3.

##### 2.3 MINFLUX tracking of Histone H2B-Halo and *Fbn2* locus dynamics

MINFLUX tracking data was acquired on a commercial microscope from Abberior Instruments GmbH similar to the setup described by Schmidt et al. (*12*). In brief, it was built around an inverted microscope body (IX83, Olympus) equipped with a 1.4 NA UPLXAPO 100X oil immersion objective (Olympus), featured a 561nm and a 640 nm continuous-wave laser for MINFLUX and confocal recordings, and was operated via the iMSPECTOR Image Acquisition and Analysis

Software (v16.3.15645, Abberior Instruments GmbH). Fluorescence emission was detected with avalanche photodiodes with the pinhole set to 0.83AU in the 580- to 630nm spectral range for 561 nm tracking experiments, and in the 650- to 720-nm spectral range for 640 nm tracking measurements. Proper alignment was confirmed daily prior to use. The tracking sequence was optimized to meet the dynamics of the samples, probing with a hexagonal pattern with diameters ranging between L = 30 - 250 nm, a dwell time of 150 µs, photon limit of 10, estimator coefficients for 20 photons (H2B tracking) or 30 (array tracking), and a background threshold of 120 kHz in the last iteration. For array tracking, the center probing position was not recorded, and the initial 561nm laser power was set to 390 µW (measured at the scanner). For H2B tracking, the initial 640nm laser power was set to 550 µW. For tracking measurements with L = 30 - 40 nm / L = 100 nm / L = 150 - 250 nm, the power was increased by a factor of 6 / 4 / 2 in the last iteration. The laser power was chosen such that the average number of photons detected in the last iteration matched the photon number of the estimator coefficients in the sequence. The raw data was exported in .msr format from iMSPECTOR (v16.3.15647, Abberior Instruments GmbH) and converted to .npy format offline (iMSPECTOR v16.3.15620, Abberior Instruments GmbH).

#### 3 Data processing

##### 3.1 MINFLUX

The MINFLUX method directly produces particle trajectories, without the intermediate of camera images. Quality control and data cleaning thus have to be done at the trajectory level, without taking recourse to movies. We apply the following processing steps:

- determine time lag and assign integer time coordinates;
- cut off the first localization (which might contain artifacts from MINFLUX initialization);
- prune background tracks (most dominant in *Fbn2* TetO array tracking, c.f. section 3.1.1);
- set a minimum trajectory length of 1000 frames;
- filter out stuck trajectories (only for U2OS H2B tracking, c.f. section 3.1.2).

While not camera-based, MINFLUX still operates in discrete time: the unit of time is one localization cycle, whose length depends on the acquisition settings (e.g. number of excitation positions). However, the `.npy` files exported from the iMSPECTOR software (section 2.3) contain localizations in the format $(I_i, x_i, y_i, t_i)$, where $I_i$ is an integer identifying which trajectory the localization $(x_i, y_i)$ belongs to and $t_i$ is the absolute time since beginning of the acquisition in seconds, i.e. a continuous time coordinate. To robustly extract the fundamental time lag $\Delta t$ (effective length of the acquisition cycle) from the continuous $t_i$ for each acquisition run, we

- compute the time steps $\delta_i \equiv t_{i+1} - t_i$ for all $i$ where $I_{i+1} = I_i$ (i.e. localizations $i+1$ and $i$ belong to the same trajectory);
- find a rough estimate
$$\widetilde{\Delta t} = \underset{\Delta t}{\operatorname{argmin}} \sum_i \left( \delta_i - \Delta t \left[ \frac{\delta_i}{\Delta t} \right] \right)^2 , \tag{1}$$
where square brackets $[\cdot]$ indicate rounding to nearest integer;
- sort the continuous time lags into bins $\left[ \left(n - \frac{1}{2}\right) \widetilde{\Delta t}, \left(n + \frac{1}{2}\right) \widetilde{\Delta t} \right)$, $n \in \mathbb{N}$ and calculate the bin averages $\bar{\delta}_n$;
- find the final estimate $\widehat{\Delta t}$ by least-squares fitting $\bar{\delta}_n = \widehat{\Delta t}\, n$.

The time lags $\widehat{\Delta t}$ obtained by this method were consistent across acquisitions for both H2B and TetO tracking, with means of $\widehat{\Delta t}_{\text{H2B}} \approx 216\,\mu\text{s}$ and $\widehat{\Delta t}_{\text{TetO}} \approx 191\,\mu\text{s}$, respectively (fig. S1). Further analysis uses these consensus time steps.

###### 3.1.1 *Fbn2* locus

Specifically for our tracking of the *Fbn2* TetO array, there are many "background" trajectories, where the system picks up and tracks spurious noise signals (fig. S2A). These background tracks have a tendency to move straight downwards (ballistic motion in the negative $y$ direction) and can thus be filtered out quite easily, as detailed below. They can occur as "pure background tracks" (no real signal) or at the beginning or end of a "true" trajectory.

Since MINFLUX trajectories are very long (tens of thousands of frames), we do not need single-frame precision in pruning background tracking. We therefore use the following simple algorithm:

- for both coordinates $(x(t), y(t))$, calculate the median value over the whole trajectory;
- find the first and last time the coordinate crosses this median, which we label $t_{\text{first}}^{x/y}$ and $t_{\text{last}}^{x/y}$, respectively;
- retain only that part of the trajectory which is between $\max\left(t_{\text{first}}^{x}, t_{\text{first}}^{y}\right)$ and $\min\left(t_{\text{last}}^{x}, t_{\text{last}}^{y}\right)$;
- omit any trajectory where that remaining part is shorter than 1000 frames.

For an example of the effect of this filtering, see fig. S2B for cutting of background trajectories and fig. S2C for subsequent length thresholding. This method successfully prunes background tracking—which is expected to behave ballistically and thus not return to previous positions—while retaining most of the "true" trajectories; which are expected to be subdiffusive and thus exhibit frequent median crossings. Note that we calculate median crossing times for the individual coordinate traces $x(t)$ and $y(t)$; these are 1D, where even diffusive processes would exhibit recurrence and thus frequent median crossings. So, while the efficiency of this pruning method does increase for more strongly subdiffusive data (as expected for chromatin and confirmed by visual inspection of trajectories, e.g. in fig. S2), it does not rely on particularly strong assumptions about the observed dynamics. It does, however, introduce a selection bias in our data set, where the retained trajectories are very likely to return to their starting point at the end of the trajectory (since both are close to the median of at least one coordinate). In the MSD plots in fig. S2 this is apparent as single-trajectory MSD curves generally sharply dropping at the end. Due to the small statistical weight of these maximal lag times (in both the calculation of ensemble MSDs and Bayesian MSD analysis) this bias is not expected to have a noticeable effect on our analysis and we choose to ignore it.

By removing short trajectories after this pruning step we achieve complete removal of any "background only" trajectories. These would otherwise leave short trajectory snippets in the pruned data set, as apparent in fig. S2B.

#### 3.1.2 H2B in U2OS

Processing of the H2B MINFLUX data followed the procedure outlined for the TetO array above, illustrated in fig. S3A–C. Background tracking is a vastly smaller problem in these data, such that the pruning step does not have a major effect (but still removes odd trajectory ends here and there); the major filtering step here is the thresholding of trajectories to be longer than 1000 frames.

However, specifically for our data from U2OS cells, upon visual inspection of the single trajectory MSD curves after background pruning, we noticed pronounced heterogeneity. A certain fraction of the trajectories was apparently stuck, exhibiting almost flat MSD curves (fig. S3C1)—these trajectories most likely correspond to dyes stuck on the surface of the coverslip and thus need to be removed. Single-trajectory MSD fits (section 4 and fig. S3F) showed a pronounced bimodality in exponent values, suggesting that indeed these are two distinct classes of trajectories (as opposed to simply exhibiting broad heterogeneity in exponent values) and providing us with a classification criterion (95% upper confidence limit for $\alpha$ smaller than 0.1, red dotted line in fig. S3F). Stuck vs. non-stuck trajectories did not exhibit particular patterns within individual nuclei (fig. S3D2), over time (fig. S3D3), or between nuclei or acquisition runs (i.e. they did not seem to stem from, e.g. dead cells, as we observed in some mESC SPT acquisitions). Comparing MINFLUX to SPT data (fig. S3E) showed good agreement between the non-stuck fraction and SPT (accounting for the MSD distortion due to motion blur in the SPT data, c.f. main text Fig. 3A), while the stuck fraction did not seem to have a counterpart in the SPT data. We therefore decided to exclude the stuck fraction from further analysis.

###### 3.1.3 H2B in mESC

Processing of H2B MINFLUX tracking in mESC followed the same procedure as in U2OS, outlined above. Contrary to U2OS, we did not find evidence for two distinct populations of trajectories in the mESC data (fig. S4F) and the two populations identified by using the same threshold as for U2OS both agreed reasonably well with SPT tracking (fig. S4E). We therefore did not remove any trajectories from the mESC data after applying the length threshold of 1000 frames (fig. S4C).

##### 3.2 SPT

#### 3.2.1 H2B

The main challenge with SPT over tens to hundreds of seconds is the fact that living cells move. The observed motion of single histones in the laboratory rest frame typically starts being dominated by this whole cell/nucleus motion over tens of seconds. As such, it as tantamount to correct for cell motion in our 2 s time lag SPT data; for consistency and to ensure absence of cell motion we treated the 100 ms time lag SPT data equivalently.

To correct for cell motion, we consider the 2-point MSD: instead of calculating the mean squared displacement for each single particle trajectory $\boldsymbol{x}(t)$, we use two particles $\boldsymbol{x}_1(t)$ and $\boldsymbol{x}_2(t)$ and compute the MSD of the relative particle position $\boldsymbol{x}_1 - \boldsymbol{x}_2$:

$$\begin{aligned}\text{MSD}_{\text{2-point}}(\Delta t) &:= \left\langle \left[(\boldsymbol{x}_1(t+\Delta t) - \boldsymbol{x}_2(t+\Delta t)) - (\boldsymbol{x}_1(t) - \boldsymbol{x}_2(t))\right]^2 \right\rangle \quad (2)\\ &= \left\langle (\boldsymbol{x}_1(t+\Delta t) - \boldsymbol{x}_1(t))^2 + (\boldsymbol{x}_2(t+\Delta t) - \boldsymbol{x}_2(t))^2 - 2\left(\boldsymbol{x}_1(t+\Delta t) - \boldsymbol{x}_1(t)\right)\left(\boldsymbol{x}_2(t+\Delta t) - \boldsymbol{x}_2(t)\right)\right\rangle \quad (3)\\ &= \text{MSD}_{x_1}(\Delta t) + \text{MSD}_{x_2}(\Delta t) - 2\text{Cov}\left(\Delta\boldsymbol{x}_1, \Delta\boldsymbol{x}_2\right) . \quad (4)\end{aligned}$$

As eq. (4) demonstrates, the 2-point MSD as defined by eq. (2) is a sum of three terms: the 1-point MSD of particle 1, the 1-point MSD of particle 2, and the covariance of displacements of these two particles. Accordingly, if we can reasonably assume the motion of particles 1 and 2 to be equivalent ($\text{MSD}_{x_1} \equiv \text{MSD}_{x_2}$) and independent ($\text{Cov}(\Delta\boldsymbol{x}_1, \Delta\boldsymbol{x}_2) = 0$), we find that

$$\text{MSD}_{\text{2-point}}(\Delta t) = \text{MSD}_{x_1}(\Delta t) + \text{MSD}_{x_2}(\Delta t) \equiv 2\text{MSD}_{\text{1-point}}(\Delta t) . \quad (5)$$

Accordingly, we can calculate a cell motion invariant single locus MSD as 0.5 times the 2-point MSD between two independent chromosomal loci.

How do we choose these two loci? In our SPT experiments we image nuclei containing multiple labelled histones, at a density low enough to allow for correct tracking, but high enough to observe as many particles as possible (usually 10 to 50) simultaneously. We can thus calculate 2-point MSDs between neighboring particles; because of the low labelling density necessary for tracking, nearest neighbor particles are usually separated by at least 1 µm, such that we can reasonably assume them to be independent, as required by eq. (4). We furthermore restrict particles to be separated by no more than 3 µm for 2-point MSD calculation, thus making our reported results even moderately robust to deformations of single nuclei—as especially mESC are prone to do—as opposed to only rigid body translation.

Calculating 2-point MSDs from every possible locus pair would be highly redundant and thereby introduce statistical artifacts. Consider, for example, three particles $\boldsymbol{x}_1(t)$, $\boldsymbol{x}_2(t)$, and $\boldsymbol{x}_3(t)$ and the associated 2-point trajectories

$$\begin{aligned}\boldsymbol{y}_{12}(t) &:= \boldsymbol{x}_1(t) - \boldsymbol{x}_2(t) , \quad (6)\\ \boldsymbol{y}_{23}(t) &:= \boldsymbol{x}_2(t) - \boldsymbol{x}_3(t) , \quad (7)\\ \boldsymbol{y}_{31}(t) &:= \boldsymbol{x}_3(t) - \boldsymbol{x}_1(t) . \quad (8)\end{aligned}$$

Clearly $\boldsymbol{y}_{12}(t) + \boldsymbol{y}_{23}(t) + \boldsymbol{y}_{31}(t) = 0$, such that one of these trajectories can always be reconstructed from the other two. To obtain a 2-point data set free of redundancy, we thus need to choose one out of the three to omit. Now, particles 1, 2, and 3 might not always be observed at the same time: due to photobleaching and intermittently missing localizations, at any given time $t$ we might or might not have a localization for particle $i$. Accordingly, we can calculate the relative position $\boldsymbol{y}_{ij}(t)$ only at times $t$ where both particles $i$ and $j$ have a valid localization. In the three particle example above, particles 1 and 2 might be visible in frames 1 through 200; while particle 3 only briefly appears in frames 87 through 132. In this case, when selecting the redundancy-free data set, we can maximize the amount of data retained by choosing $y_{23}$ or $y_{31}$ for omission (either containing only 46 localizations) and making sure to retain $y_{12}$—which contains 200 valid localizations.

Generalizing to $N > 3$ particles, we address this need for a maximal, redundancy-free 2-point data set as follows:

- from trajectories $\boldsymbol{x}_1(t), \ldots, \boldsymbol{x}_N(t)$ in any given nucleus, compute all pairwise trajectories $\boldsymbol{y}_{ij}(t) := \boldsymbol{x}_i(t) - \boldsymbol{x}_j(t)$.
- for each pairwise trajectory $\boldsymbol{y}_{ij}(t)$, count the number $F_{ij}$ of valid frames (i.e. the number of frames where both particles $i$ and $j$ are localized).
- set $F_{ij} = 0$ if either $F_{ij} < 20$ (too little overlap), or $\langle|\boldsymbol{x}_i(t) - \boldsymbol{x}_j(t)|\rangle_t > 3\,\mu\text{m}$ (particles too far apart).
- construct an undirected graph $G$ with $N$ nodes (corresponding to the individual particles) and edge weights $F_{ij}$.
- find the maximum spanning tree of $G$, defined as the set of edges with maximum total weight that does not contain any cycles. Technically we achieve this by constructing a graph $G'$ with edge weights $-F_{ij}$ and using `scipy.sparse.csgraph.minimum_spanning_tree()` to find the minimum spanning tree of $G'$; which is equivalent to the maximum spanning tree of $G$.

The edges in the maximum spanning tree of $G$ constitute an (almost) maximal, redundancy-free 2-point data set.

To see that absence of cycles in the graph $G$ is equivalent to absence of redundancy in the set of 2-point trajectories $Y$, note that $y_{ij} = -y_{ji}$ and "redundancy" here refers to the existence of any index set $I \subset \{(i,j) \mid i,j = 1,\ldots,N\}$ such that $y_{ij} \in Y\, \forall (i,j) \in I$ and

$$\sum_{(i,j)\in I} y_{ij} \equiv 0 . \quad (9)$$

Since $y_{ij} = x_i - x_j$, the above summation condition holds true if and only if the index set $I$ is such that for any $y_{\cdot i}$ (containing a $-x_i$ term) there is a corresponding $y_{i\cdot}$ (containing a $+x_i$ term), implying that $I$ can be ordered as

$$I = \{(a,b), (b,c), \dots, (m,n), (n,a)\} \ , \tag{10}$$

where the $a, ..., n$ do not necessarily have to be distinct. Index sets of this form are exactly the cycles in $G$; cycles in the graph thus correspond exactly to redundancies in the data set.

Maximality of the data set does not strictly hold, as it is easy to construct scenarios in which our procedure would not produce all possible two-point trajectories free of redundancy. Under realistic conditions, however, these "missed trajectories" will be few, short, and statistically equivalent to the rest of the data (i.e. we do not introduce bias by omitting them). Thus, while theoretically our method could still be improved to obtain the truly maximum amount of data, we do not pursue this optimization here.

###### 3.2.2 *Fbn2* locus

To correct for cell motion in the SPT data from the *Fbn2* locus, we also calculated 2-point MSDs, in this case between the integrated TetO array and non-specifically labelled CTCF. Processing followed closely that for H2B SPT, i.e. we omitted particle pairs that either were separated (on average) by more than 3 µm or had fewer than 20 valid localizations. In contrast to H2B SPT, each pair now consisted of two different particles (TetO and CTCF), such that there was no redundancy issue. We thus used all 2-point trajectories fulfilling the mentioned conditions on length and separation.

##### 3.3 SRLCI

SRLCI data for clones C36 and F1M were used as previously published (*3*) without additional processing. We used only $x$ and $y$ coordinates for compatibility with the 2D trajectories of MINFLUX and SPT, omitting the axial position of the loci. However, because the tracking was originally in 3D, we do not loose particles due to diffusion out of the focal plane, which would be a concern for "truly 2D" tracking data at these time scales. We calculated the 2-point MSD between the two tracked loci according to eq. (2), thus obtaining a cell motion invariant observable. As for cell motion corrected SPT data, we multiplied this 2-point MSD by 0.5 for plotting in main text Figs. 3D and 4D, thus ensuring to study comparable observables.

Due to the genomic proximity (505 kb) of the two loci, the independence condition leading to eq. (5) is violated at long times, where the MSD plateaus to the limiting value $2\left\langle R^2 \right\rangle$. At short times (before the crossover in MSD, in these data around 30 min), one would expect the two loci to not "feel" their genomic connection, thus moving independently and restoring validity of eq. (5). A possible interpretation of the mismatch between SPT and SRLCI data in Figs. 3D and 4D is that even at the 20 s shortest time scale in the SRLCI data, that is not true; suggesting that even at these short time scales the two loci can "feel" their connection. This is an interesting observation warranting further study.

##### 3.4 Relative mobility calculation in Fig. 4 insets

To provide a simple summary of the effects of our drug treatments, we averaged the log-ratio of treatment over control MSD over all available time scales. Specifically:

$$\text{relative mobility(treatment)} := \frac{\displaystyle\sum_{\substack{\text{technique}\in\\ \{\text{MINFLUX,SPT,SRLCI}\}}} \int_{\log \Delta t_{\text{min}}^{\text{technique}}}^{\log \Delta t_{\text{max}}^{\text{technique}}} \left(\log \text{MSD}_{\text{treatment}}^{\text{technique}} - \log \text{MSD}_{\text{control}}^{\text{technique}}\right) \mathrm{d}\log\Delta t}{\displaystyle\sum_{\substack{\text{technique}\in\\ \{\text{MINFLUX,SPT,SRLCI}\}}} \left(\log \Delta t_{\text{max}}^{\text{technique}} - \log \Delta t_{\text{min}}^{\text{technique}}\right)} \ . \tag{11}$$

We average in log-space (integration with respect to $\log \Delta t$ instead of $\Delta t$) to achieve equal weighting of all available time scales.

##### 3.5 Summary

- MINFLUX data was cleaned and cut to a minimum length of 1000 localizations per trajectory. For plotting in main figures, we truncate MINFLUX MSDs at $\Delta t = 1\,\text{s}$.
- SPT data was converted to 2-point data set to remove cell motion; for consistency with MINFLUX we plot 0.5 times 2-point MSD. Trajectories were cut to a minimum length of 20 localizations per trajectory. For plotting in main figures, we truncate SPT MSDs to 100 frames, corresponding to 10 s for SPT (100 ms) and 200 s for SPT (2 s), respectively.

- SRLCI data was used without additional processing, save for removing the axial coordinate ($z$). Similarly to SPT, we plot 0.5 times 2-point MSD. For plotting in main figures, we truncate SRLCI MSDs to 300 frames, corresponding to 6000 s or 1 h 40 min.

Untruncated, single-trajectory MSD plots for all conditions, stratified by replicate, are provided in figs. S6 to S15.

#### 4 Bayesian MSD fitting

For MSD fitting we employed the methodology developed by Gabriele, Brandão, Grosse-Holz *et al.* (*3*). Briefly, any MSD curve together with a suitable stationarity condition (stationarity for e.g. 2-point MSD of tethered loci; increment stationarity for e.g. free (sub-)diffusion) uniquely defines a Gaussian process and thus a probability distribution $P\left[x(t) \mid \text{MSD}(\Delta t)\right]$ over trajectories $x(t)$ (square brackets indicating that $P$ is a path measure over trajectories $x(t)$ instead of a probability distribution over values $x \equiv x(t)$ for some fixed $t$). Given a trajectory $x(t)$ (or multiple trajectories), this probability distribution can now be used as a likelihood function over MSD($\Delta t$). Upon introducing a prior over possible MSD functions that enforces some parametric form (e.g. $\text{MSD}(\Delta t) = \Gamma \Delta t^{\alpha}$), this likelihood function over the functional space of MSD functions reduces to an unnormalized posterior density over a finite-dimensional parameter space (the space of $(\Gamma, \alpha)$ in the power law example). We can then numerically find the maximum *a posteriori* (MAP) parameters, i.e. those that maximize the posterior. We report these MAP parameters as fit results.

Having the full (unnormalized) posterior density at our disposal also allows for further analysis, e.g. calculating credible intervals. To that end, we numerically trace the profile likelihood in parameter space until hitting a threshold set by the desired credibility level (95% by default)(*3*). We use this credible bound on single-trajectory exponents $\alpha$ to identify stuck trajectories in U2OS (section 3.1.2 and fig. S3F).

##### 4.1 Fitting MINFLUX data

In our MINFLUX data, ensemble MSDs seem to exhibit a “kink” at the shortest time scales, apparent e.g. in main text Fig. 3C. Since we could reproduce this effect in our MINFLUX simulations (fig. S5), we concluded that this is a technical artifact and should be excluded from fitting. To achieve this exclusion, we subsampled our MINFLUX trajectories by a factor 4 before applying the Bayesian MSD fitting, thus effectively cutting lag times $< 4\Delta t$ from the fit.

Furthermore, MINFLUX trajectories can be very long, with the number of localizations $F$ per trajectory numbering in the tens of thousands. This poses a computational problem for the Bayesian MSD fitting, which relies on inversion of a covariance matrix of dimension $F \times F$ and thus scales poorly with trajectory length. To overcome this computational issue, we applied a hierarchical subsampling approach:

- define a suitable maximum trajectory length, we choose $T = 200$ time steps;
- cut the original trajectory into snippets of length $T$;
- ensure that each snippet starts with a valid localization by cutting off possible invalid localizations from the beginning;
- save the snippets and associated time lag $\Delta t$ (time between consecutive localizations);
- assemble a subsampled (coarse-grained) trajectory consisting of only the first localization of each snippet; the associated time lag $\Delta t_{\text{cg}}$ is given by $\Delta t_{\text{cg}} = T\Delta t$;
- iterate the cutting and coarse-graining steps on this subsampled trajectory, saving the generated snippets for each time lag, until the resulting coarse-grained trajectory is shorter than the maximum trajectory length $T$.

With this approach we convert a data set of MINFLUX trajectories (individual long trajectories) into a data set of shorter snippets at different time scales, thus still covering the whole dynamics range of MINFLUX. We then jointly fit this whole data set, taking into account the different lag times for each set of snippets. Intuitively, this approach corresponds to cutting the MINFLUX MSD curve into segments of $T = 200$ data points each and then fitting them jointly, instead of fitting the whole curve at once. This allows us to fit long MINFLUX trajectories with acceptable computational effort.

##### 4.2 Motion blur (and localization error)

Real particle tracking data contains technical artifacts that need to be considered in analysis. Most prominently, we can never determine the location of a particle exactly, leading to finite localization accuracy $\sigma > 0$. This is usually assumed to be uncorrelated and Gaussian (i.e. white noise), thus resulting in an additive term $+2d\sigma^2$ in the MSD ($d$ being space dimension). Another, less frequently discussed source of bias in MSD curves is *motion blur*: we can usually not determine a particle's location at a single time point, but have to integrate over some short time window (exposure time for camera-based tracking or acquisition cycle for MINFLUX). If this integration time is comparable to the lag time between localizations, this will lead to correlations in the observed motion and a corresponding bias in the MSD. In this section we calculate the effects of motion blur and localization error, showing how the *observed* MSD $\psi_Y(\Delta t)$ depends on *true* MSD $\psi_X(\Delta t)$ of the observed particle, integration time (formulated as an *shutter function* $\zeta(\tau)$), and localization accuracy $\sigma$.

Let us assume that we image the stochastic process $X(t)$ at discrete times $\{t_n\}$, thus obtaining the collection of random variables

$$Y_n \equiv \int_0^{\Delta T} \mathrm{d}\theta\, \zeta(\theta) X(t_n - \theta) + \sigma \xi_n \,, \tag{12}$$

where $\Delta T \equiv \min_{m,n|m\neq n} |t_m - t_n|$ is the minimum separation between time points, $\zeta : [0, \Delta T] \to \mathbb{R}_+$ is the *shutter function*[*] satisfying $\int_0^{\Delta T} \mathrm{d}\theta\, \zeta(\theta) = 1$, $\sigma$ is the standard deviation of localization error, and $\xi_n$ are uncorrelated standard normal random variables.

Note that we did not assume even spacing of the $\{t_n\}$, although this is overwhelmingly the most common imaging modality. However, experimental data may have gaps (missing frames), which makes the spacing uneven (though still integral multiples of $\Delta T$). For sake of generality we then simply assume fully unevenly spaced $\{t_n\}$. We then, however, have to take care of a few details; the following list should be taken as footnotes to the calculations below.

We define $\tau_{mn} \equiv t_m - t_n$ and note that

- $\tau_{km} + \tau_{mn} = \tau_{kn}$, since the $\tau$ are signed.
- We require $\tau_{mn} \neq 0 \,\forall m \neq n$, i.e. differently indexed times should be different.
- For $n \neq m$, the integration domains for $Y_m$ and $Y_n$ in eq. (12) should not overlap, otherwise the localization error term would pick up correlations. This motivates the choice of $\Delta T = \min_{m,n|m\neq n} |\tau_{mn}|$.
- Make sure to distinguish continuous time variables and discrete indices. This will be especially important in the following study of increments, where clearly the "increment process" should be defined with a lag*time*, not a lag*index*; but this lagtime will only be allowed to take values from subsets of $\{\tau_{mn}\}$.
- Below we will write Kronecker $\delta$'s over the time indices; note that with the above requirement that $\tau_{mn} \neq 0 \,\forall m \neq n$ these can be expressed purely in terms of $\tau_{mn}$, since $\delta_{mn} = 1 \Leftrightarrow \tau_{mn} = 0$.

##### 4.2.1 Exact calculation of imaging artifacts for general MSD

We define the increments of $Y$ as

$$\left(\Delta^{\tau_{mn}} Y\right)_n \equiv Y_m - Y_n = \int_0^{\Delta T} \mathrm{d}\theta\, \zeta(\theta) \Big[X(t_m - \theta) - X(t_n - \theta)\Big] + \sigma\left(\xi_m - \xi_n\right) \,. \tag{13}$$

These increments are correlated according to

$$C_Y^{\tau_{kl},\tau_{mn}}(\tau_{ln}) := \Big\langle \left(\Delta^{\tau_{kl}} Y\right)_l \left(\Delta^{\tau_{mn}} Y\right)_n \Big\rangle_{\mathsf{c}} \tag{14}$$

$$\equiv \Big\langle \left(Y_k - Y_l\right)\left(Y_m - Y_n\right) \Big\rangle_{\mathsf{c}} \tag{15}$$

$$= \int_0^{\Delta T} \mathrm{d}\theta\, \mathrm{d}\theta'\, \zeta(\theta)\zeta(\theta') \Big\langle \Big[X(t_k - \theta) - X(t_l - \theta)\Big]\Big[X(t_m - \theta') - X(t_n - \theta')\Big] \Big\rangle_{\mathsf{c}} \tag{16}$$

$$+ \sigma^2 \Big\langle \left(\xi_k - \xi_l\right)\left(\xi_m - \xi_n\right) \Big\rangle_{\mathsf{c}}$$

$$= \int_0^{\Delta T} \mathrm{d}\theta\, \mathrm{d}\theta'\, \zeta(\theta)\zeta(\theta')\, C_X^{\tau_{kl},\tau_{mn}}(\tau_{ln} + \theta' - \theta) + \sigma^2\left(\delta_{km} - \delta_{lm} - \delta_{kn} + \delta_{ln}\right) \,, \tag{17}$$

[*]Here we are assuming that light emission by the fluorophore is continuous in time. However, considering single photons arriving as inhomogeneous Poisson process with time dependent rate $\lambda(\tau)$ reproduces this treatment in expectation, with the shutter function $\zeta(\tau) \propto \lambda(\tau)$. We omit the discrete treatment here for brevity.

where $C_X$ is the analogous increment correlation of the true process $X(t)$, satisfying

$$C_X^{\tau_1,\tau_2}(\Delta t) := \Big\langle \Big[X(t+\Delta t+\tau_1) - X(t+\Delta t)\Big]\Big[X(t+\tau_2) - X(t)\Big]\Big\rangle_{\text{c}} \tag{18}$$

$$= \Big\langle X(t+\Delta t+\tau_1)X(t+\tau_2) - X(t+\Delta t)X(t+\tau_2) - X(t+\Delta t+\tau_1)X(t) + X(t+\Delta t)X(t)\Big\rangle_{\text{c}} \tag{19}$$

$$\begin{aligned} = \frac{1}{2}\Big\langle & 2X(t+\Delta t+\tau_1)X(t+\tau_2) && - X(t+\Delta t+\tau_1)^2 && - X(t+\tau_2)^2 \\ & -2X(t+\Delta t)X(t+\tau_2) && + X(t+\Delta t)^2 && + X(t+\tau_2)^2 \\ & -2X(t+\Delta t+\tau_1)X(t) && + X(t+\Delta t+\tau_1)^2 && + X(t)^2 \\ & +2X(t+\Delta t)X(t) && - X(t+\Delta t)^2 && - X(t)^2 \Big\rangle_{\text{c}} \end{aligned} \tag{20}$$

$$= \frac{1}{2}\Big[\psi_X(\Delta t - \tau_2) + \psi_X(\Delta t + \tau_1) - \psi_X(\Delta t + (\tau_1 - \tau_2)) - \psi_X(\Delta t)\Big]. \tag{21}$$

Note also that

$$C_X^{\Delta t,\Delta t}(0) = \Big\langle \Big[X(t+\Delta t) - X(t)\Big]\Big[X(t+\Delta t) - X(t)\Big]\Big\rangle_{\text{c}} = \psi_X(\Delta t)\,, \tag{22}$$

with the equivalent relations holding for $C_Y$. Using these relations and transforming the integration variables as

$$\vartheta \equiv \theta' - \theta \in [-\Delta T, \Delta T]\ , \tag{23}$$

$$\bar\theta \equiv \frac{\theta' + \theta}{2} \in \left[\frac{|\vartheta|}{2}, \Delta T - \frac{|\vartheta|}{2}\right]\ , \tag{24}$$

we can rewrite eq. (17) in terms of the MSDs $\psi_X$ and $\psi_Y$, for $\Delta t > 0$:

$$\psi_Y(\Delta t) = C_Y^{\Delta t,\Delta t}(0) \tag{25}$$

$$= 2\sigma^2 + \int_{-\Delta T}^{\Delta T} \mathrm{d}\vartheta\, Z(\vartheta) C_X^{\Delta t,\Delta t}(\vartheta) \tag{26}$$

$$= 2\sigma^2 + \frac{1}{2}\int_{-\Delta T}^{\Delta T} \mathrm{d}\vartheta\, Z(\vartheta)\Big[\psi_X(\vartheta+\Delta t) + \psi_X(\vartheta - \Delta t) - 2\psi_X(\vartheta)\Big]\,, \tag{27}$$

where

$$Z(\vartheta) \equiv \int_{\frac{|\vartheta|}{2}}^{\Delta T - \frac{|\vartheta|}{2}} \mathrm{d}\bar\theta\, \zeta\left(\bar\theta + \frac{\vartheta}{2}\right)\zeta\left(\bar\theta - \frac{\vartheta}{2}\right)\,. \tag{28}$$

Note that $Z(-\vartheta) = Z(\vartheta)$ and $\int_{-\Delta T}^{\Delta T} \mathrm{d}\vartheta\, Z(\vartheta) = 1$ by the normalization of $\zeta(\theta)$. Furthermore, by definition $\psi_X(-\Delta t) = \psi_X(\Delta t)$, so we can exploit these parity symmetries and the symmetric $\vartheta$-integration domain to write

$$\int_{-\Delta T}^{\Delta T} \mathrm{d}\vartheta\, Z(\vartheta)\psi(\vartheta - \Delta t) = \int_{-\Delta T}^{\Delta T} \mathrm{d}\vartheta\, Z(\vartheta)\psi(\Delta t - \vartheta) = \int_{-\Delta T}^{\Delta T} \mathrm{d}\vartheta\, Z(\vartheta)\psi(\Delta t + \vartheta)\,, \tag{29}$$

such that finally we find

$$\psi_Y(\Delta t) = 2\sigma^2 + \int_{-\Delta T}^{\Delta T} \mathrm{d}\vartheta\, Z(\vartheta)\psi_X(\Delta t + \vartheta) - \int_{-\Delta T}^{\Delta T} \mathrm{d}\vartheta\, Z(\vartheta)\psi_X(\vartheta)\,. \tag{30}$$

Below we will study the individual contributions to this expression in more detail. For intuition, note that

- The first term is the standard additive constant due to localization error
- The second term is a "washed-out" version of the original MSD $\psi_X$. Since $Z(\vartheta)$ is normalized, a first approximation for this term is simply $\int_{-\Delta T}^{\Delta T} \mathrm{d}\vartheta\, Z(\vartheta)\psi_X(\Delta t + \vartheta) = \psi_X(\Delta t)$.
- The third term does not depend on $\Delta t$ and thus presents another—negative—additive constant. Note that it depends only on the MSD at $\vartheta \in [-\Delta T, \Delta T]$, i.e. lag times below a single frame.

###### 4.2.2 Solution for powerlaw MSD and "all-or-nothing" illumination

To investigate the behavior of eq. (30) in more detail, let us consider MSDs of the shape $\psi_X(\Delta t) = \Gamma\,|\Delta t|^\alpha$, $\alpha \in [0,2)$ and shutter functions of the shape

$$\zeta(\tau) = \frac{1}{f\Delta T}\Theta\left(f\Delta T - \tau\right) \equiv \begin{cases} \frac{1}{f\Delta T} & \tau \in [0, f\Delta T]\ , \\ 0 & \text{else}\,, \end{cases} \tag{31}$$

with $f \in [0,1]$ the *fractional exposure time*. Direct calculation then shows that

$$Z(\vartheta) = \frac{1}{f\Delta T}\left(1 - \frac{|\vartheta|}{f\Delta T}\right)\Theta\left(f\Delta T - |\vartheta|\right) \qquad \forall f > 0\,, \tag{32}$$

while for $f = 0$ (ideal stroboscopic illumination) we find $Z(\vartheta) = \delta(\vartheta)$.

Having found an expression for $Z(\vartheta)$ then allows us to calculate the integrals in eq. (30):

$$\int_{-\Delta T}^{\Delta T} \mathrm{d}\vartheta\, Z(\vartheta)\psi_X(\Delta t + \vartheta) = \Gamma \int_{-f\Delta T}^{f\Delta T} \frac{\mathrm{d}\vartheta}{f\Delta T}\left(1 - \frac{|\vartheta|}{f\Delta T}\right)|\Delta t + \vartheta|^\alpha \tag{33}$$

$$\text{let } \varphi \equiv \frac{f\Delta T}{\Delta t} \text{ and substitute } x \equiv \frac{\vartheta}{f\Delta T} \tag{34}$$

$$= \Gamma\,|\Delta t|^\alpha \int_{-1}^{1} \mathrm{d}x\ (1 - |x|)\,|1 + \varphi x|^\alpha \tag{35}$$

$$= \Gamma\,|\Delta t|^\alpha\, \frac{|1+\varphi|^{\alpha+2} + |1-\varphi|^{\alpha+2} - 2}{\varphi^2(\alpha+1)(\alpha+2)} \tag{36}$$

$$\equiv \psi_X(\Delta t) b(\Delta t, f, \alpha)\,. \tag{37}$$

Clearly the integral in the third term of eq. (30) is just the same expression, evaluated at $\Delta t = 0$. It can straightforwardly be calculated from eq. (33), or obtained as the $\varphi \to \infty$ asymptote of eq. (36). Taking the latter route here gives

$$\int_{-\Delta T}^{\Delta T} \mathrm{d}\vartheta\, Z(\vartheta)\psi_X(\vartheta) = \Gamma\,|\Delta t|^\alpha\, \frac{2\,|\varphi|^\alpha}{(\alpha+1)(\alpha+2)} \tag{38}$$

$$= \frac{2\Gamma\,|f\Delta T|^\alpha}{(\alpha+1)(\alpha+2)} \tag{39}$$

$$\equiv 2B\left(\Gamma, \alpha, f\right)\ . \tag{40}$$

Finally, eq. (30)—for powerlaw MSD $\psi_X$—reads

$$\psi_Y(\Delta t) = \psi_X(\Delta t) b(\Delta t) - 2B + 2\sigma^2\,, \tag{41}$$

with

$$b(\Delta t) := \frac{|1+\varphi|^{\alpha+2} + |1-\varphi|^{\alpha+2} - 2}{\varphi^2(\alpha+1)(\alpha+2)} \qquad \left(\varphi \equiv \frac{f\Delta T}{\Delta t}\right)\,, \tag{42}$$

$$B := \frac{\psi_X(f\Delta T)}{(\alpha+1)(\alpha+2)}\,. \tag{43}$$

We employ eq. (41) in the Bayesian MSD fitting scheme to account for motion blur.

###### 4.2.3 Motion blur for low exponents

Motion blur becomes extremely strong for low exponents. Consider the extreme (and toy) example of white noise, $\alpha = 0$, where each localization, at any time scale, is sampled uniformly from a standard normal Gaussian distribution. Applying any finite integration time to this process, one would average an infinite sample from that Gaussian and thus obtain the analytical mean of the distribution, $0$. White noise, imaged with any finite integration time, is a constant.

Similarly, but not quite as extreme, when imaging highly subdiffusive processes with a finite integration time, much of the fluctuations at short time scales average out, such that the observed motion seems significantly more static (less mobile) than the true motion (as observable in e.g. main text Fig. 3).

We make this intuition quantitative by studying eq. (41), assuming negligible localization error $\sigma = 0$ for the time being. Equation (41) holds for powerlaw MSDs $\psi_X$ which, using the notation of eq. (42), we write as

$$\psi_X(\varphi) = \Gamma\varphi^{-\alpha} \,. \tag{44}$$

Note that this implies that we measure time in multiples of the integration time $f\Delta T$; we furthermore assume $\varphi^{-1} = \frac{\Delta t}{f\Delta T} > 1$. Now, using eqs. (42) and (43), eq. (41) reads

$$\psi_Y(\varphi) = \Gamma\varphi^{-\alpha}\frac{(1+\varphi)^{\alpha+2} + (1-\varphi)^{\alpha+2} - 2}{\varphi^2(\alpha+1)(\alpha+2)} - \frac{2\Gamma}{(\alpha+1)(\alpha+2)} \,, \tag{45}$$

$$= \frac{2\Gamma}{(\alpha+1)(\alpha-1)}\left[\frac{1}{2}\left(\varphi^{-1}+1\right)^{\alpha+2} + \frac{1}{2}\left(\varphi^{-1}-1\right)^{\alpha+2} - \left(\varphi^{-1}\right)^{\alpha+2} - 1\right] \,, \tag{46}$$

$$= \alpha\Gamma\left[\frac{1}{2}\left(\varphi^{-1}-1\right)\log\left(\varphi^{-1}-1\right) - \varphi^{-2}\log\varphi^{-1} + \frac{1}{2}\left(\varphi^{-1}+1\right)\log\left(\varphi^{-1}+1\right)\right] + \mathcal{O}\left(\alpha^2\right) \,. \tag{47}$$

This result is quite interesting: for low exponents $\alpha$, the shape of the observed MSD (the term in square brackets) becomes independent of $\alpha$! Instead, the observed MSD $\psi_Y(\Delta t)$ is completely dominated by motion blur and adopts a universal shape, only scaled by a prefactor $\alpha\Gamma$; it thus becomes impossible to distinguish $\alpha$ and $\Gamma$ experimentally.

Note that the above expansion for small $\alpha$ holds at constant $\varphi^{-1} = \frac{\Delta t}{f\Delta T}$; meanwhile, at constant $\alpha$, of course we recover $\psi_Y(\Delta t) \to \psi_X(\Delta t)$ as $\Delta t \to \infty$ (in the sense of $\frac{\psi_Y}{\psi_X} \to 1$; can be shown by expanding eq. (42) to second order in $\varphi$ and noting that $B$ is constant, such that $\frac{B}{\psi_X} \to 0$ as $\Delta t \to \infty$). So, while the shape of the MSD close to the integration time becomes universal for small $\alpha$, for times much larger than the integration time the MSD always recovers its true behavior, as one would expect.

Fortunately, for exponent values of $\alpha \approx 0.3$, as reported in this work, this collapse of the observed MSD is not a major issue yet. It does, however, have implications for the numerics of our fitting procedure. To obtain meaningful results from e.g. the single trajectory fits we use for identification of stuck particles in section 3.1.2, we parameterize the pertinent powerlaw fits in terms of $(\log\alpha\Gamma, \alpha)$, instead of the more obvious pair $(\log\Gamma, \alpha)$. This ensures that if the fit reaches the universal regime, we still get meaningful results for the prefactor $\alpha\Gamma$ and an upper bound on $\alpha$, instead of an arbitrarily chosen pair $(\Gamma, \alpha)$ from the solutions of $\alpha\Gamma = \text{const}$.

##### 4.3 Fit results

Numerical results of our fits are provided in table S2. Visualization on top of the experimental data is shown in figs. S16 to S19.

#### 5 Simulations

##### 5.1 First-passage times

The paths of fractional Brownian motion with an exponent $\alpha$ have fractal dimension $d_f = \frac{2}{\alpha}$. If $d_f$ is larger than the space dimension $d$, an infinitely long fBm path will densely fill the available space, eventually reaching any point in said space. Put another way, an fBm searcher will find a point-like target, if $\alpha < \frac{2}{d} = 0.66$ (in 3D space). The pertinent search time distribution has a powerlaw tail and diverging mean, unless the search is confined to a finite volume (Guerin et al., Nature 2016); we therefore use the median instead of mean as characteristic time scale of the search process. The median has the added advantage of being robustly estimable from simulation data as the time until 50% of sampled trajectories hit the target; we do not need to sample the long tail of the distribution exhaustively.

The question in this section is the following: what is the median search time (first passage time) $\tau$ for a $d$-dimensional fractional Brownian motion with MSD$(\Delta t) = \Gamma_d\Delta t^\alpha$, $\alpha < \frac{2}{d}$ to find a point-like target initially a distance $X$ away? Note that this analytical problem is fully determined by the constants $\alpha$ and $d$ (dimensionless), $X$ (with units of length), and $\Gamma_d$ (with units of length$^2$/time$^\alpha$). Any expression for $\tau$ can thus be made up of only these constants. By dimensional analysis, this has to take the form

$$\tau = q_d(\alpha)\left(\frac{X^2}{\Gamma_d}\right)^{\frac{1}{\alpha}} \,, \tag{48}$$

with $q_d(\alpha)$ a numerical prefactor dependent on the dimensionless $\alpha$. Here we are predominantly interested in $d = 3$ and thus introduce the shorthand $q(\alpha) := q_3(\alpha)$, i.e. we will drop the explicit reference to space dimension.

Equation (48) can also be understood by the following scaling argument: fBm with $\alpha < \frac{2}{d}$ explores its local surroundings densely; so the search time should be given (up to a numerical prefactor) by the time over which the MSD covers the distance $X$. We write

$$\tau \propto \mathsf{MSD}^{-1}(X^2) = \left(\frac{X^2}{\Gamma_d}\right)^{\frac{1}{\alpha}} , \tag{49}$$

where $\mathsf{MSD}^{-1}(\cdot)$ refers to the inverse function of $\mathsf{MSD}(\cdot)$, given explicitly on the right hand side. The only difference from eq. (48) is that there we introduced an explicit name for the proportionality constant; which we can of course do here as well. Writing $\mathsf{FPT}(X)$ instead of $\tau$, we can give an operationally very straight-forward definition:

$$q_d(\alpha) := \frac{\mathsf{FPT}(X)}{\mathsf{MSD}^{-1}(X^2)} , \tag{50}$$

which we use below.

To determine the numerical prefactor $q(\alpha)$, we resort to simulations. We simulated 1000 fractional Brownian motions of length $10^8$ for $\alpha \in \{0.2, 0.3, 0.4, 0.5, 0.6\}$ (and $\Gamma = 1$, i.e. $\mathsf{MSD}(\Delta t) = d\Gamma\Delta t^\alpha = 3\Delta t^\alpha$ in simulation units) using a custom implementation of the Davies-Harte algorithm (*13*–*17*). First passage times were then calculated as the number of steps required for the trajectory to reach a spherical target volume of radius $r$ a distance $x$ away for 8 logarithmically spaced values of $r$ between $\sqrt{3}$ and $5\sqrt{3}$ and 15 logarithmically spaced values of $x$ between $2\sqrt{3}$ and $2^{3.5}\sqrt{3}$. We estimated search time distributions using the Kaplan-Meier estimator implemented in `scikit-survival` (v0.23.1) (fig. S20A) and calculated the median search time for any parameter set where more than 50% of simulations found the target (fig. S20B). In line with our physical expectations, median search time seemed to depend predominantly on the distance to the closest point of the target volume ("effective target distance"; red lines in fig. S20B), especially for the lower exponent values. For $\alpha = 0.5, 0.6$, i.e. closer to the critical $\alpha = 0.66$, we observed an additional decrease in search time for larger targets at the same effective target distance. For small targets, our discretized simulation trajectories have a considerable risk of stepping "through" the target volume, i.e. missing search events. We therefore restricted our subsequent analysis to targets with $r > 3$. As expected from fig. S20B, we found that median FPTs collapsed quite well when plotted as a function of effective target distance $X$. Dividing by the scaling expectation $\mathsf{MSD}^{-1}(X^2)$ showed a reasonable degree of convergence for $X > 10$ (fig. S20C). We averaged all simulation results in this regime ($X > 10$ and $r > 3$) to obtain estimates for $q(\alpha)$ (fig. S20D). Note that $q(\alpha)$ describes only the median search time; the associated distribution generally spans multiple orders of magnitude (fig. S20A).

The $q(\alpha)$ estimated from our fBm simulations now allowed us to calculate FPT scalings for exemplary power law MSDs. To define a suitable family, we considered our power law fit to U2OS data ($\alpha = 0.29$, $\Gamma = 0.0101\,\mu m^2/s^{0.29}$; table S2) and a previously reported parametrization of a Rouse model ($\alpha = 0.5$, $\Gamma = 0.0024\,\mu m^2/s^{0.5}$)(*3*). These two power laws intersect at $\Delta t_{\mathsf{ref}} \approx 1100\,\mathrm{s}$, $X_{\mathsf{ref}} \approx 280\,\mathrm{nm}$, allowing us to define a family

$$\mathsf{MSD}_\alpha(\Delta t) := 2dX_{\mathsf{ref}}^2 \left(\frac{\Delta t}{\Delta t_{\mathsf{ref}}}\right)^\alpha \tag{51}$$

of MSDs with different exponent values $\alpha$. We add a prefactor 2 to model the dynamics of two independent genomic loci searching each other (c.f. eq. (4)). The associated median first passage times are then given by

$$\mathsf{FPT}_\alpha(X) = q(\alpha)\Delta t_{\mathsf{ref}} \left(\frac{X^2}{2dX_{\mathsf{ref}}^2}\right)^{\frac{1}{\alpha}} \tag{52}$$

and plotted in main text Fig. 4E.

To give rough genomic length scales associated with the distances over which we plot Fig. 4E, we use an independent calibration procedure: We(*3*) previously reported a root-mean-square separation of 547 nm between two loci 505 kb apart (in clone C65 where the CTCF sites were deleted; making these loci more representative for distances of average genomic loci), while a single nucleosome contains about 200 bp and has a diameter of 11 nm. Interpolating these two data points with a power law gives the conversion

$$X_{\mathrm{nm}} = 0.7833\,\mathrm{nm/bp}^{0.4987} S_{\mathrm{bp}}^{0.4987} . \tag{53}$$

#### 5.2 Motion blur in MINFLUX

To determine whether trajectories from MINFLUX tracking can be adequately modeled by motion blur and localization error alone, we simulated 150 fractional Brownian motions for the equivalent of 11000 MINFLUX cycles for alpha values 0.1, 0.15, 0.2, 0.25, and 0.3. We then simulated MINFLUX tracking on these trajectories varying a number of MINFLUX parameters using a custom MINFLUX simulation implementation (REF). To simulate MINFLUX, an ideal donut geometry was used as the excitation pattern with the following functional form(*18*)

$$I_{donut}(r,z) = 4\mathrm{e}\ln 2\frac{r^2}{\text{fwhm}^2}\exp\left(-4\ln 2\frac{r^2}{\text{fwhm}^2}\right)\frac{1}{\sqrt{2\pi}}\exp\left(\frac{-z^2}{2\sigma_z^2}\right) \tag{54}$$

To match the real experiment to the best of our knowledge, we used fwhm$_r$ = 360 nm, $\sigma_r$ = 350 nm, a 20 µs detection gate with a 5 µs lag time between detections and a 5 µs computation gate. Additionally, probing was performed using a hexagonal beam pattern with a center probing position corresponding to a total cycle time of 180 µs. The tracking algorithm implementation was chosen to match the real experiment to the best of our knowledge with the following parameters. We used a background emission rate of 15 000 Hz, a minimum threshold to register a localization of 10 photons, a background threshold of 80 000 Hz, a maximum dark time of 3 s, and stickiness set to 4. Since observed photon emission rates depended on the mobility of the particle, tracking was simulated for 3 relative emission rate values of 0.5, 1, and 2 $s^{-1}U^{-1}$ where U is in arbitrary units of excitation intensity. The value which made the photon emission rate max cross the photon emission threshold was selected. The modified least mean square estimator from Balzarotti *et al.* 2017 (S51) (*18*) was used with $\beta = [1, 1]$.

To determine whether motion blur and localization error alone match outputs from MINFLUX, L values of 25, 50, 100, 150, and 200 nm were used. To test the effects of different MSD prefactors, each fractional Brownian motion sample was scaled by logarithmicaly increasing values between $\sqrt{10}$ and $\sim 20$. A range of scales which approximately produced MSDs similar to what was measured in the paper was selected for display fig. S5. As for L values of 150 or higher the resulting MSD was above the theoretical ”motion-blur only” MSD, we determined that motion blur and localization error could be used to match MSDs from MINFLUX tracking with these parameters.

#### 6 Supplementary Figures

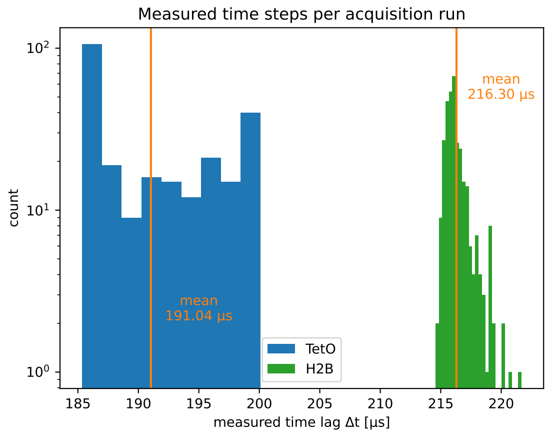

**Fig. S1:** Time steps measured from MINFLUX acquisition runs for *Fbn2* TetO array (blue) and H2B (green) tracking, with respective means (orange).

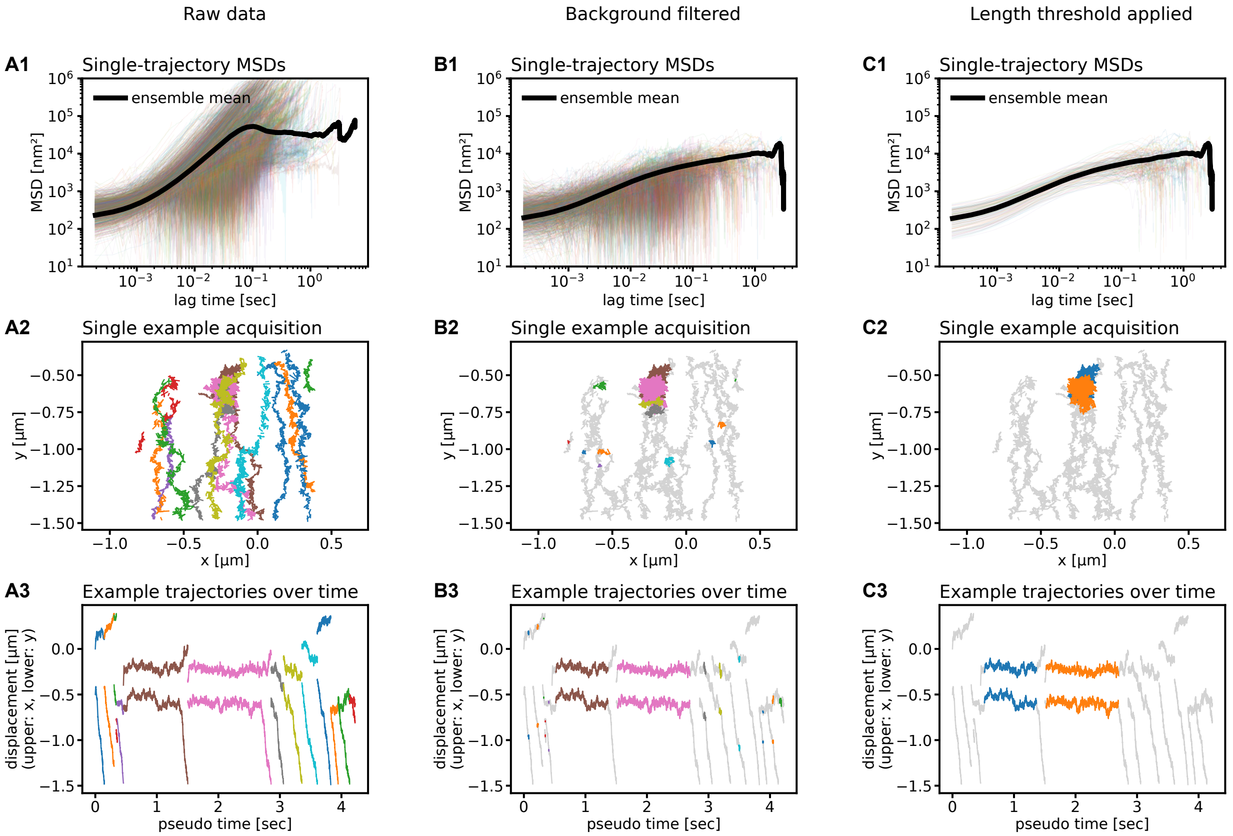

**Fig. S2:** Overview over data processing for array MINFLUX tracking. (A) The raw data contain many background traces, which need to be removed. (B) The filtering approach described in section 3.1.1 helps cutting most of the background traces, leaving only short snippets. (C) Short snippets are removed by thresholding trajectory length (1000 frames).

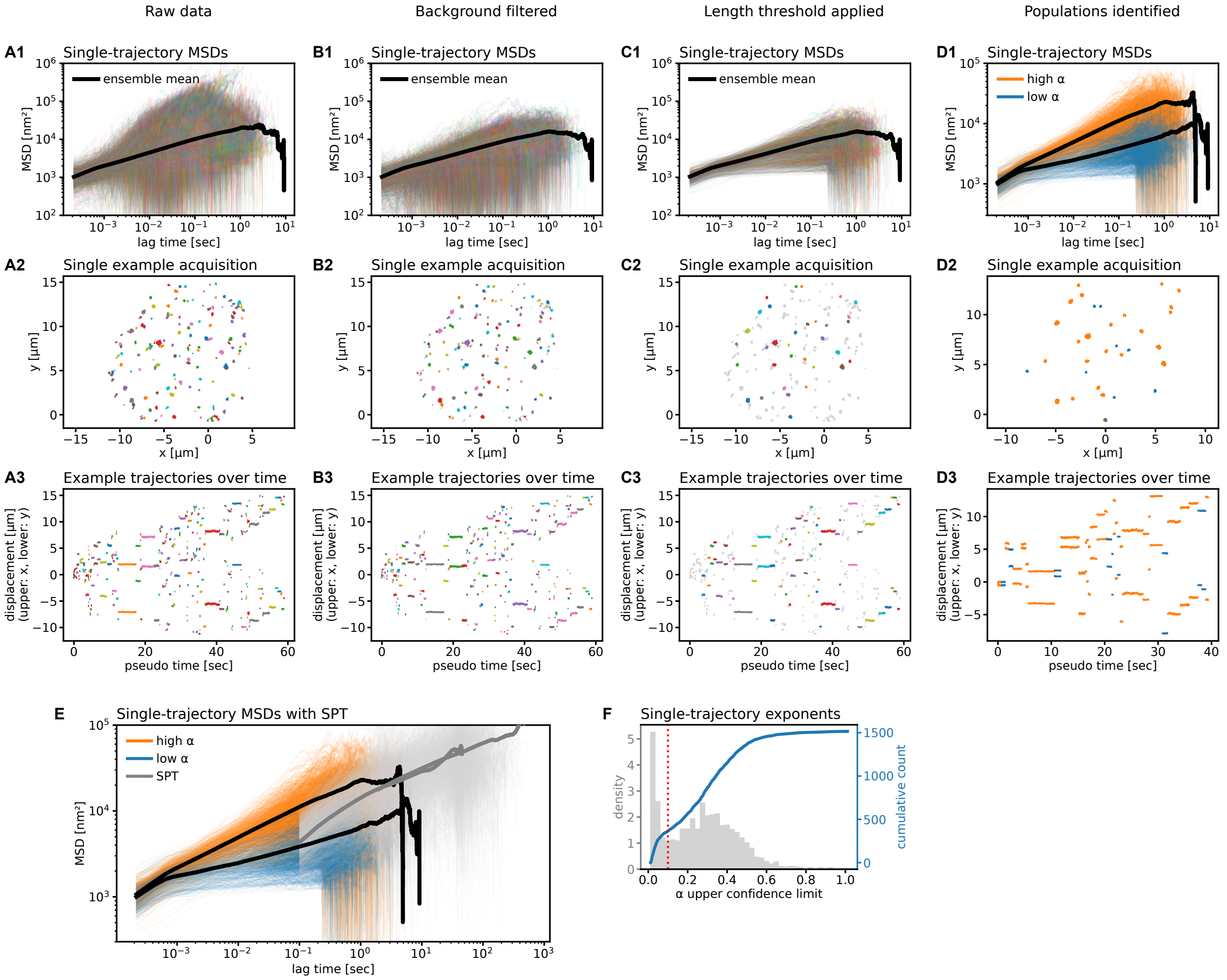

**Fig. S3:** Overview over data processing for H2B MINFLUX tracking in U2OS. (A) Raw data contains essentially no background traces. (B) Accordingly, applying background filtering does not remove a lot of data. (C) Thresholding trajectory length removes many short trajectories. Note that the ensemble MSD (black line in C1) remains virtually unaffected. (D) Identifying a stuck fraction in U2OS (see also fig. S6 and (F)). (E) Only the "non-stuck" fraction (orange) of our U2OS data seems to be represented in SPT (grey; remember that U2OS SPT slopes down at early times due to motion blur). (F) Histogram and CDF of 95% upper credible bound on single-trajectory exponent $\alpha$, exhibiting clearly bimodal behavior.

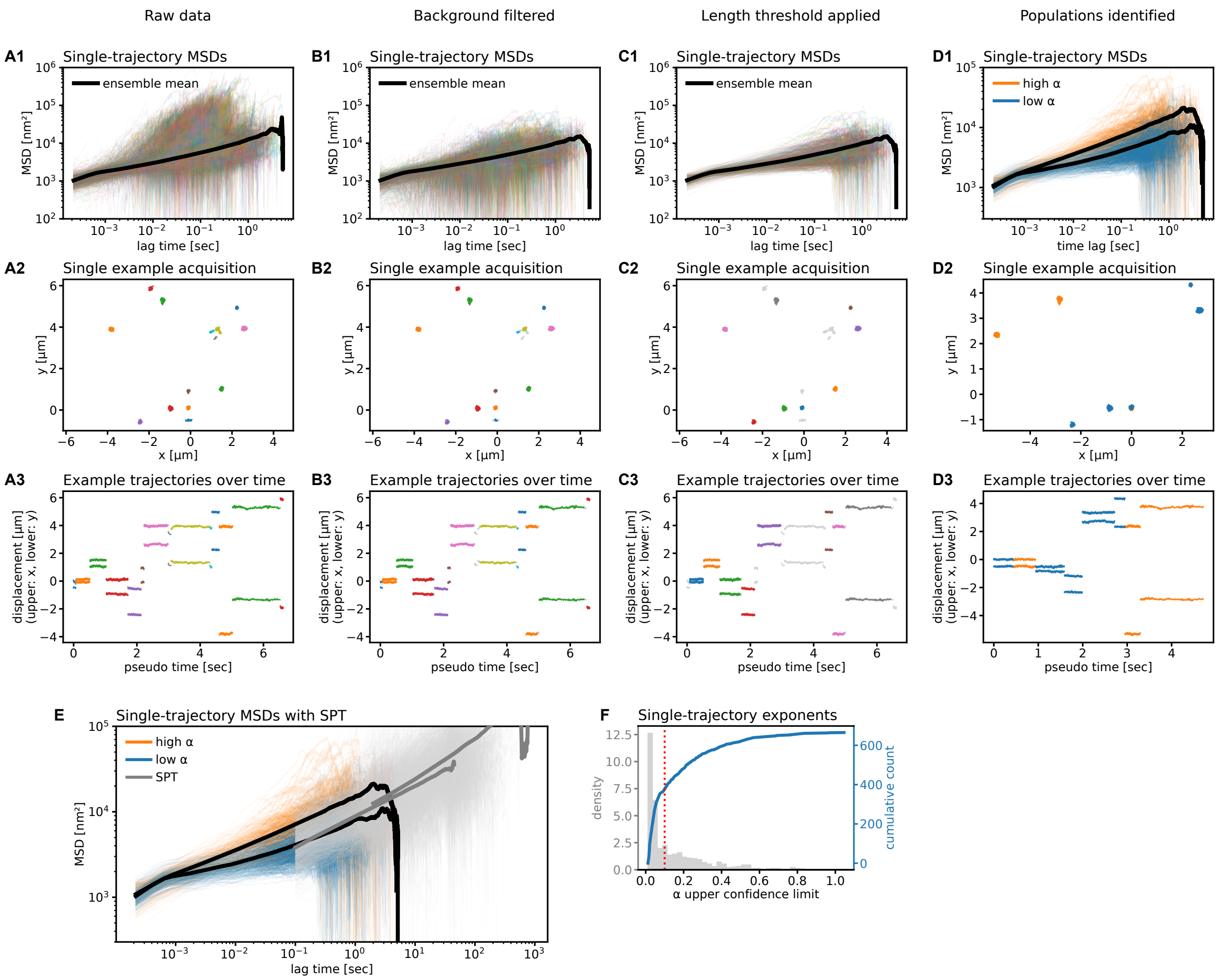

**Fig. S4:** Overview over data processing for H2B MINFLUX tracking in mESC. (A) to (D) as in fig. S3. For mESC there is no visually “stuck” fraction (as for U2OS); for consistency we still follow the same analysis as for U2OS. (E) Upon splitting mESC MINFLUX data into high and low exponent according to the procedure used for U2OS, the corresponding (mESC) SPT data seems to line up more closely with the “low exponent” fraction. However, single-trajectory MSDs (thin lines) show overlap with both fractions; upon accounting for the possibility of some motion blur (thus pushing the SPT MSD still a bit higher at early times), we conclude that our (complete) MINFLUX data set is consistent with SPT for mESC. (F) Exponent upper credible limit does not show bimodality (as opposed to U2OS, fig. S3). We therefore do *not* split the data set for further processing.

**A** Motion blur simulations

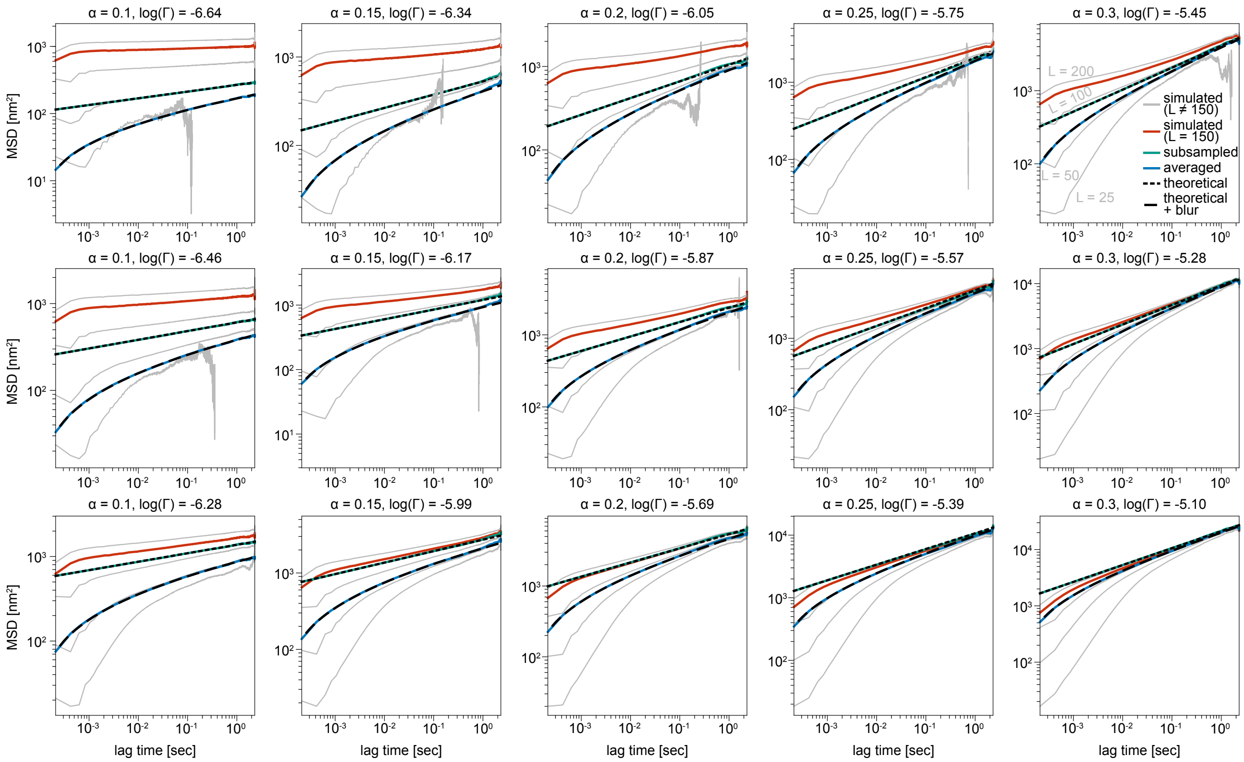

**Fig. S5:** Simulations of motion blurring in MINFLUX show L = 150 nm can be modeled by motion blurring and localization error. (A) Simulations of MINFLUX tracking of fractional Brownian motions with the given MSD exponent $\alpha$ and MSD prefactor $\Gamma$ shown above the plot. Raw trajectory MSDs are shown in teal and blue with corresponding theoretical MSDs shown with black dashed lines. MINFLUX MSDs from L values other than L = 150 nm are shown in gray, and the MSD for L = 150 nm is shown in red.

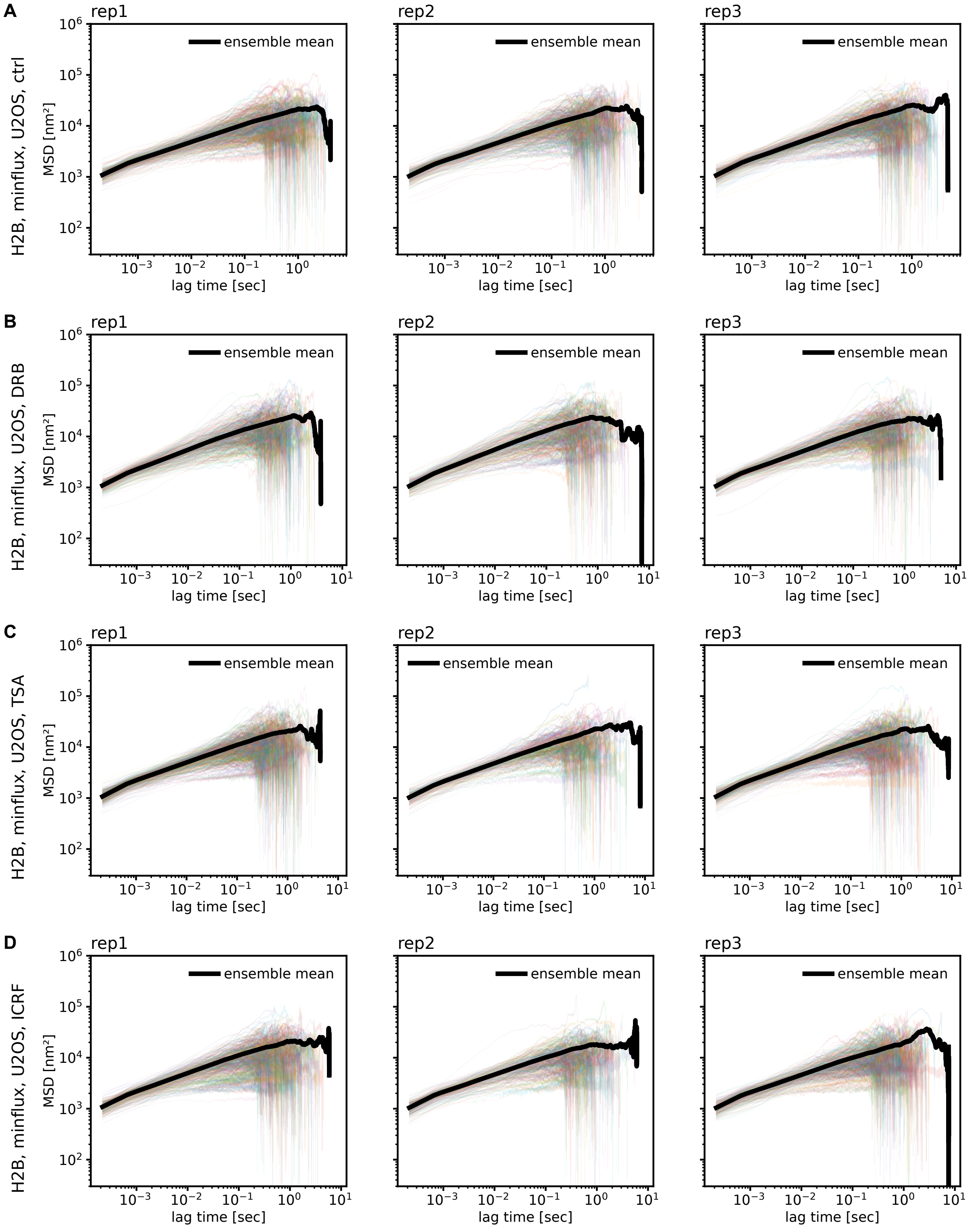

**Fig. S6:** Single trajectory MSDs for MINFLUX tracking of H2B-Halo in U2OS, stratified by biological replicate. Thin colored lines are MSDs of single trajectories, thick black is ensemble MSD. (A) control condition (DMSO). (B) DRB treatment (transcription inhibition). (C) TSA treatment (histone hyperacetylation). (D) ICRF treatment (inhibition of topoisomerase II).

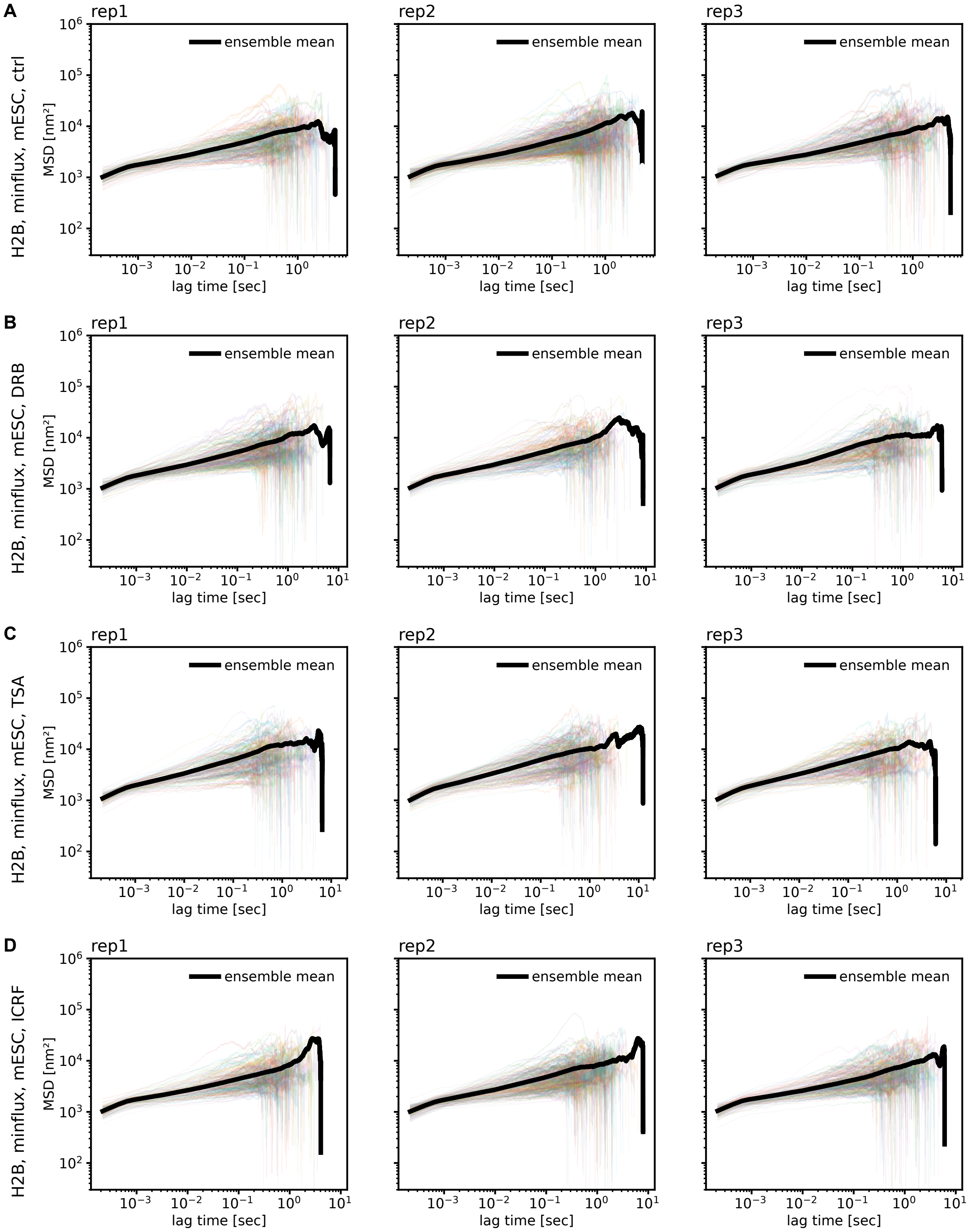

**Fig. S7:** Single trajectory MSDs for MINFLUX tracking of H2B-Halo in mESC, stratified by biological replicate. Thin colored lines are MSDs of single trajectories, thick black is ensemble MSD. (A) control condition (DMSO). (B) DRB treatment (transcription inhibition). (C) TSA treatment (histone hyperacetylation). (D) ICRF treatment (inhibition of topoisomerase II).

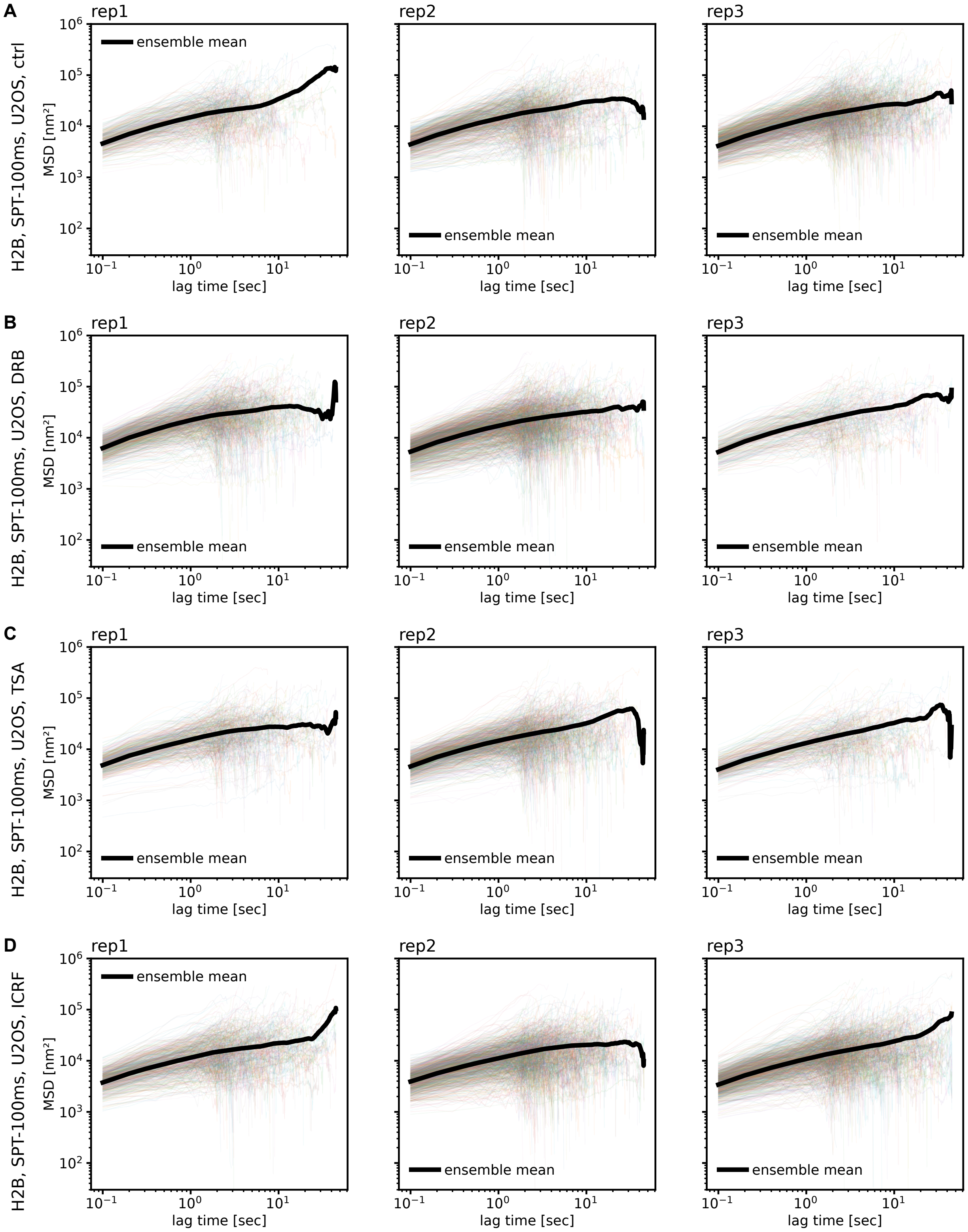

**Fig. S8:** Single trajectory MSDs for SPT tracking (100 ms lag time) of H2B-Halo in U2OS, stratified by biological replicate. Thin colored lines are MSDs of single trajectories, thick black is ensemble MSD. (A) control condition (DMSO). (B) DRB treatment (transcription inhibition). (C) TSA treatment (histone hyperacetylation). (D) ICRF treatment (inhibition of topoisomerase II).

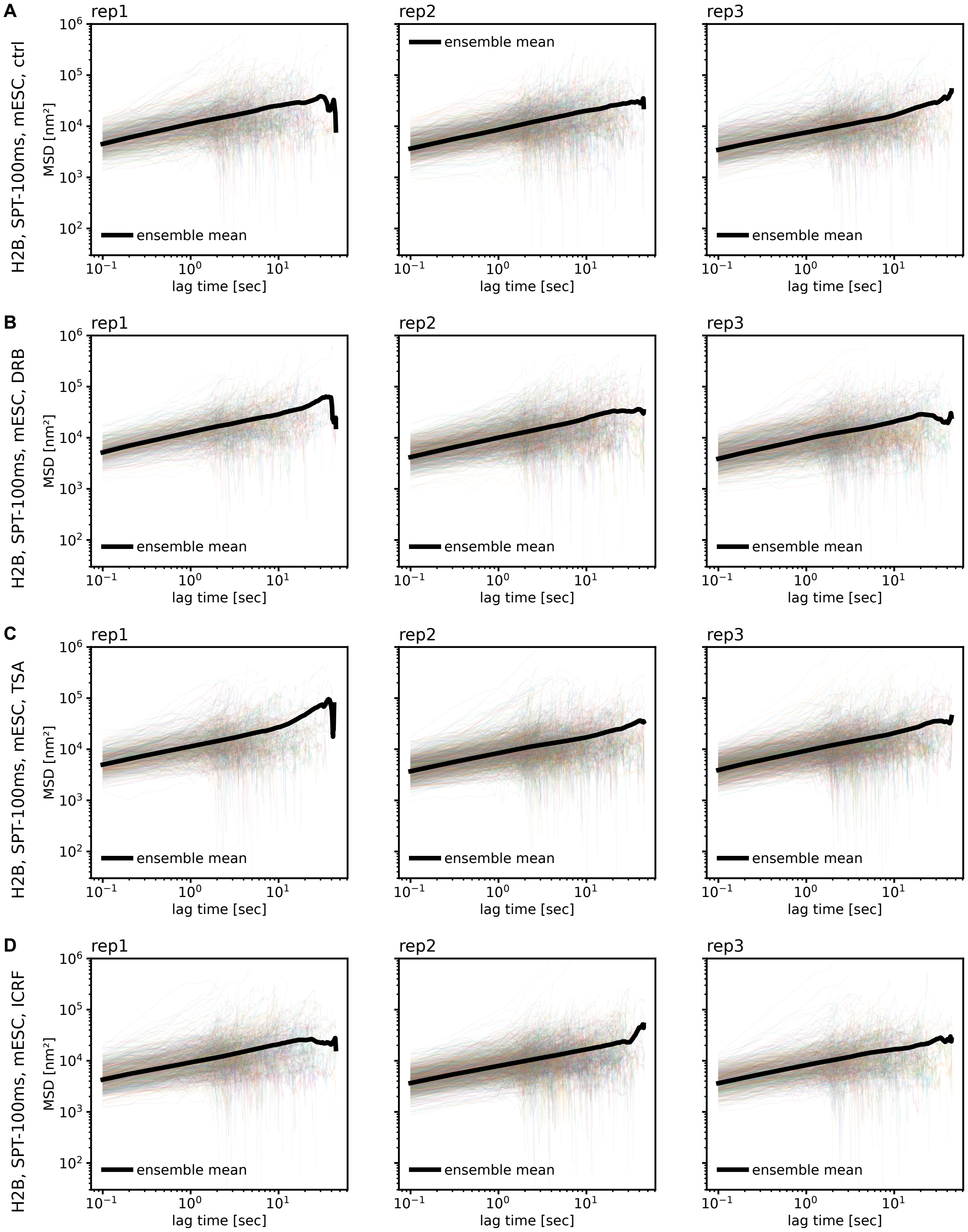

**Fig. S9:** Single trajectory MSDs for SPT tracking (100 ms lag time) of H2B-Halo in mESC, stratified by biological replicate. Thin colored lines are MSDs of single trajectories, thick black is ensemble MSD. (A) control condition (DMSO). (B) DRB treatment (transcription inhibition). (C) TSA treatment (histone hyperacetylation). (D) ICRF treatment (inhibition of topoisomerase II).

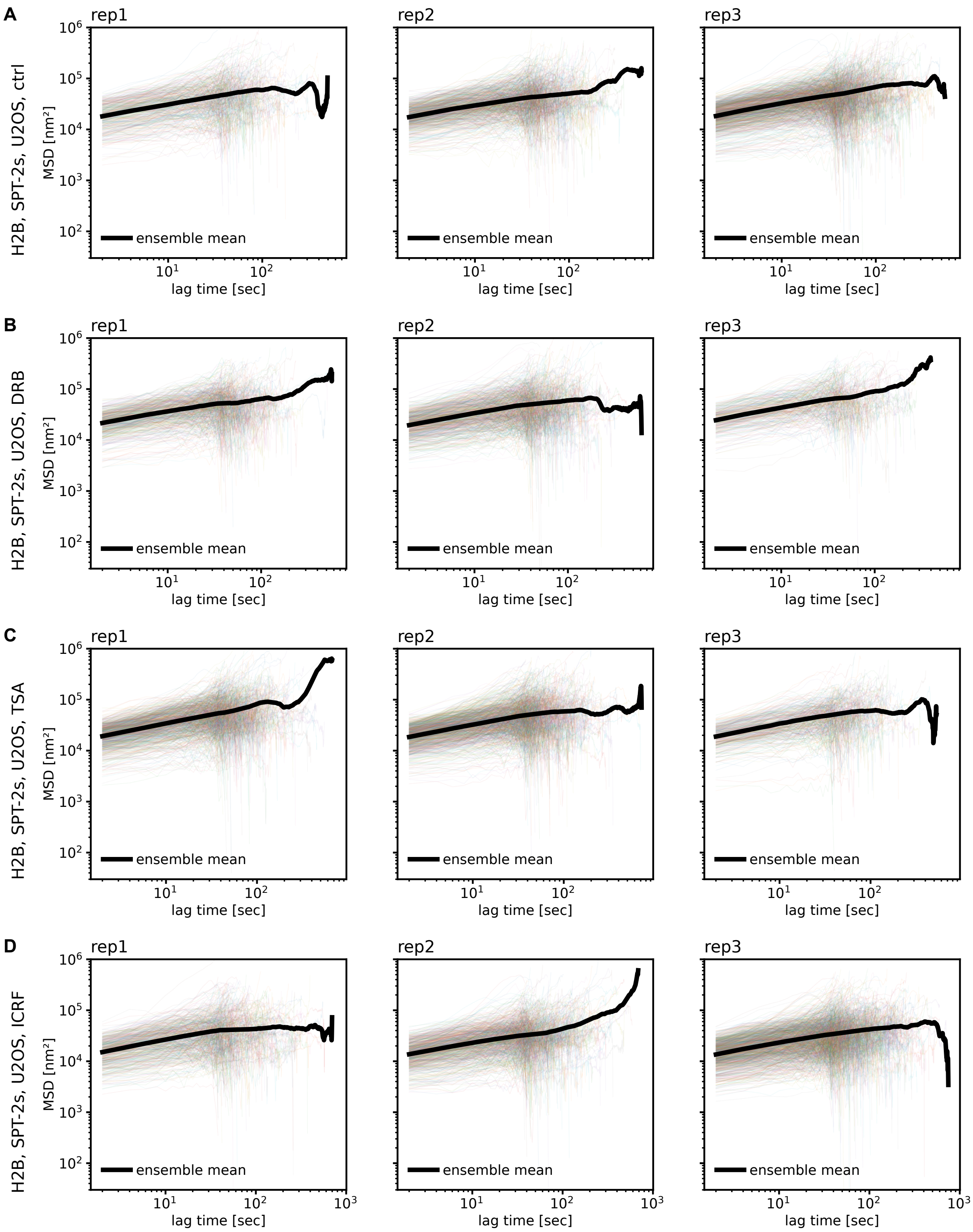

**Fig. S10:** Single trajectory MSDs for SPT tracking (2 s lag time) of H2B-Halo in U2OS, stratified by biological replicate. Thin colored lines are MSDs of single trajectories, thick black is ensemble MSD. (A) control condition (DMSO). (B) DRB treatment (transcription inhibition). (C) TSA treatment (histone hyperacetylation). (D) ICRF treatment (inhibition of topoisomerase II).

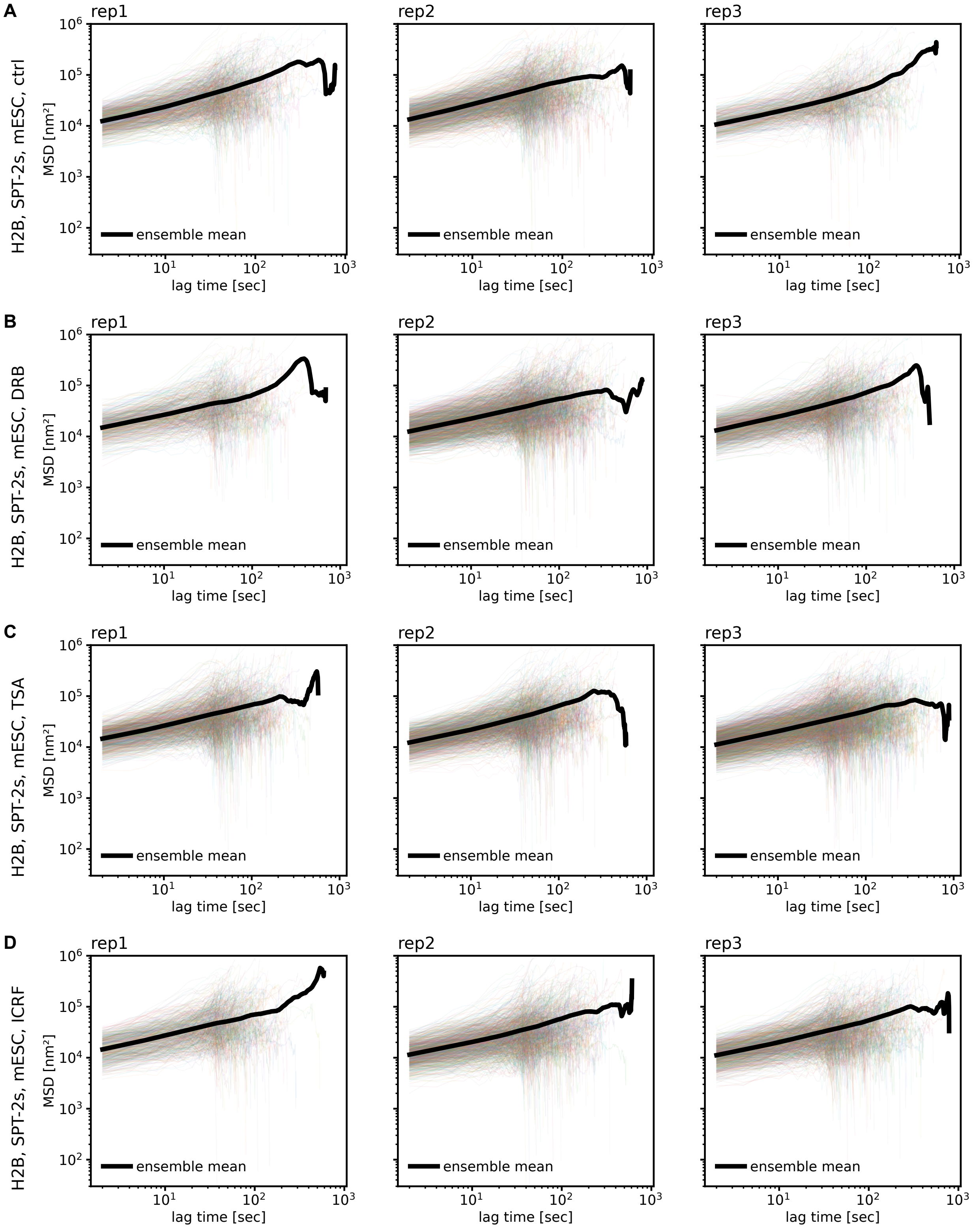

**Fig. S11:** Single trajectory MSDs for SPT tracking (2 s lag time) of H2B-Halo in mESC, stratified by biological replicate. Thin colored lines are MSDs of single trajectories, thick black is ensemble MSD. (A) control condition (DMSO). (B) DRB treatment (transcription inhibition). (C) TSA treatment (histone hyperacetylation). (D) ICRF treatment (inhibition of topoisomerase II).

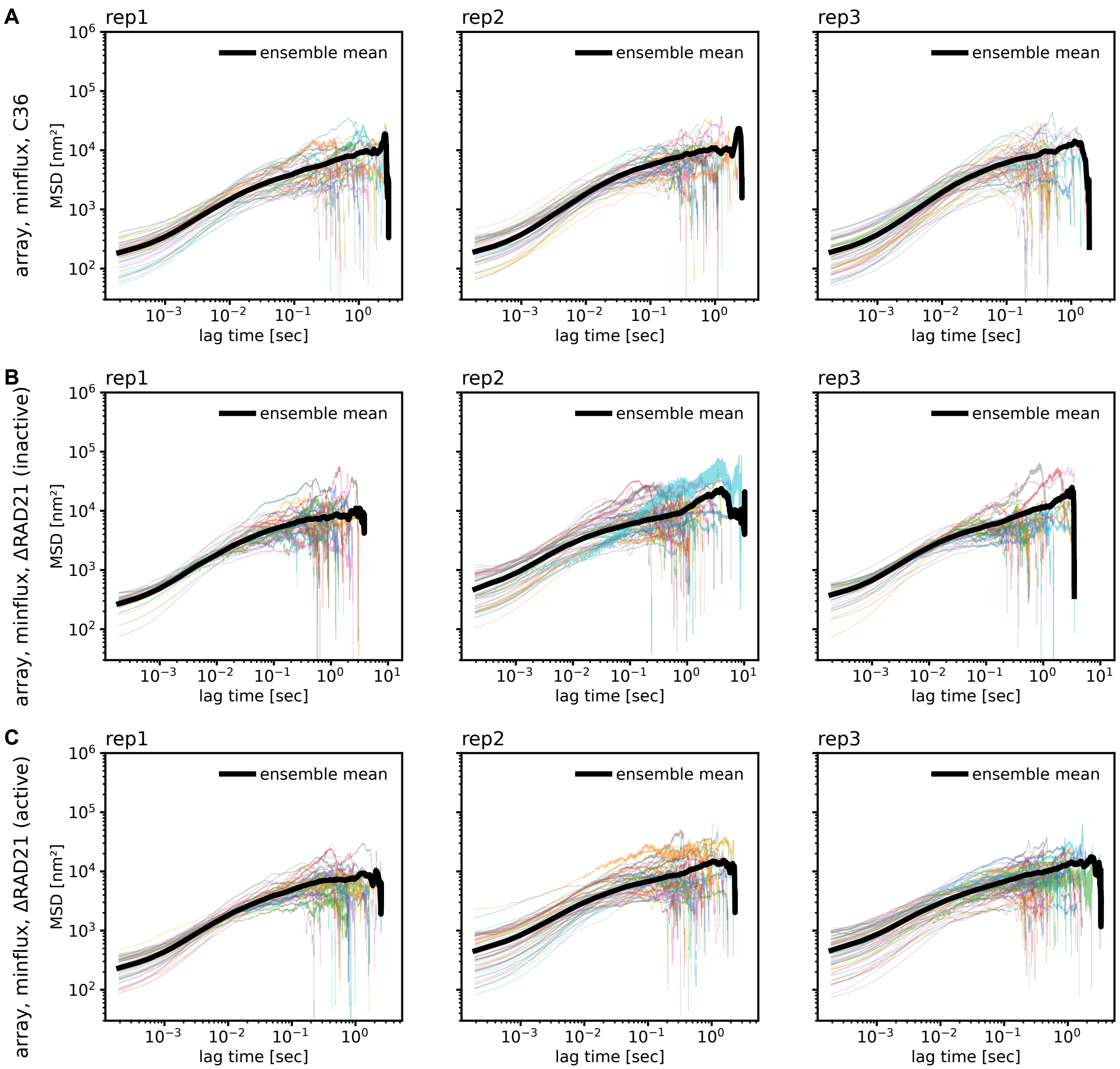

**Fig. S12:** Single trajectory MSDs for MINFLUX tracking of the *Fbn2* TetO array, stratified by biological replicate. Thin colored lines are MSDs of single trajectories, thick black is ensemble MSD. (A) control condition (DMSO). (B) DRB treatment (transcription inhibition). (C) TSA treatment (histone hyperacetylation). (D) ICRF treatment (inhibition of topoisomerase II).

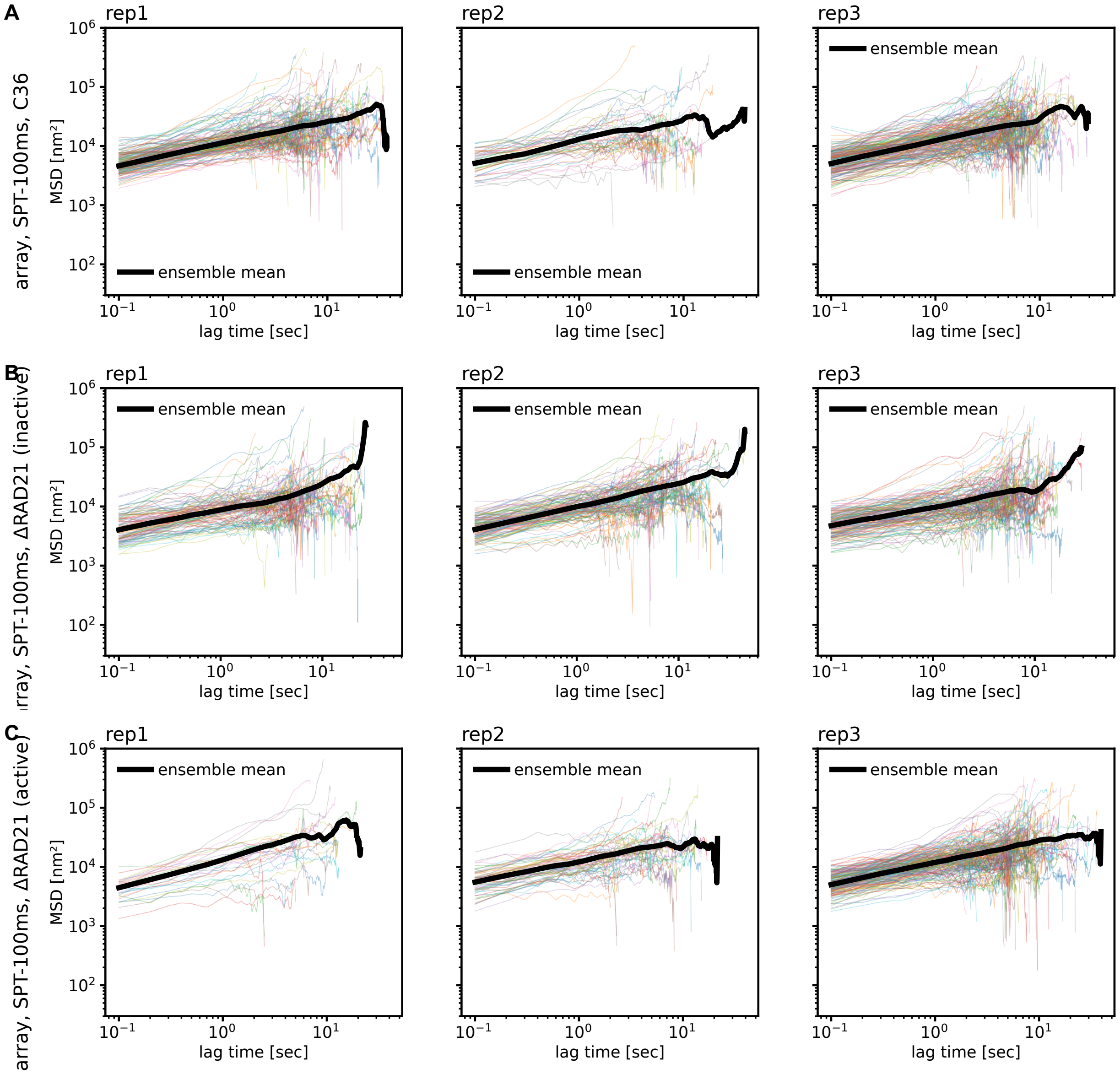

**Fig. S13:** Single trajectory MSDs for SPT tracking (100 ms lag time) of the *Fbn2* TetO array, stratified by biological replicate. Thin colored lines are MSDs of single trajectories, thick black is ensemble MSD. (A) control condition (DMSO). (B) DRB treatment (transcription inhibition). (C) TSA treatment (histone hyperacetylation). (D) ICRF treatment (inhibition of topoisomerase II).

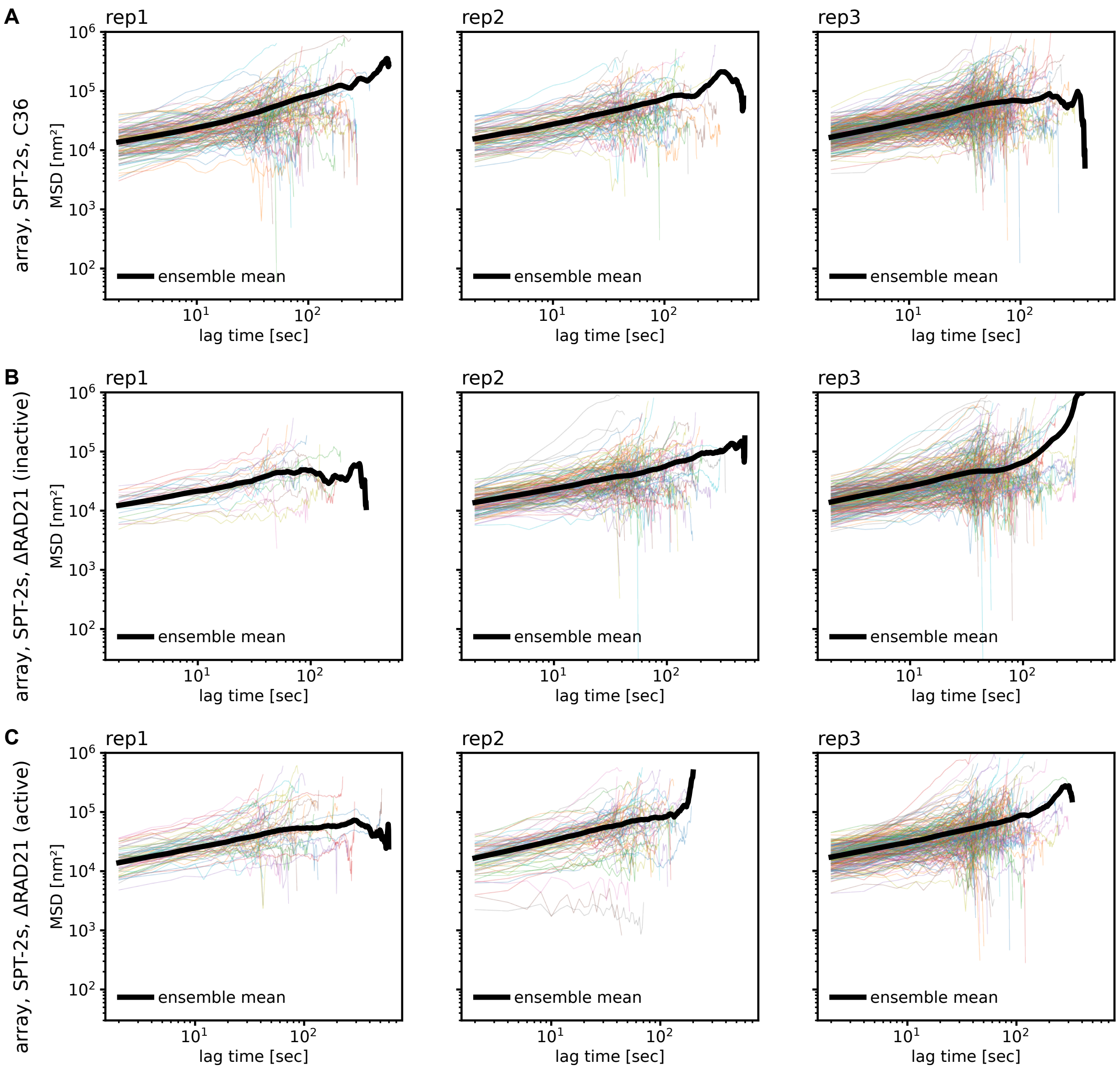

**Fig. S14:** Single trajectory MSDs for SPT tracking (2 s lag time) of the *Fbn2* TetO array, stratified by biological replicate. Thin colored lines are MSDs of single trajectories, thick black is ensemble MSD. (A) control condition (DMSO). (B) DRB treatment (transcription inhibition). (C) TSA treatment (histone hyperacetylation). (D) ICRF treatment (inhibition of topoisomerase II).

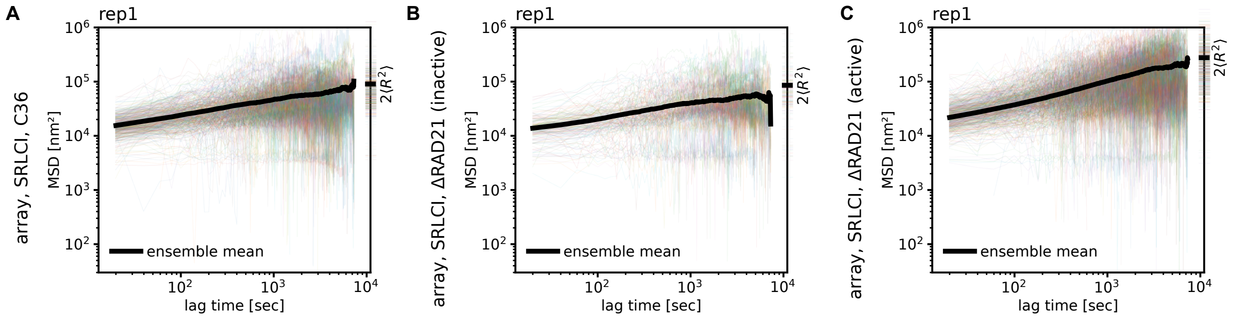

**Fig. S15:** Single trajectory MSDs for SRLCI tracking of the *Fbn2* TetO array. Biological replicates are naturally combined, since data was acquired over multiple weeks. Thin colored lines are MSDs of single trajectories, thick black is ensemble MSD. (A) control condition (DMSO). (B) DRB treatment (transcription inhibition). (C) TSA treatment (histone hyperacetylation). (D) ICRF treatment (inhibition of topoisomerase II).

treatment: None (DMSO)

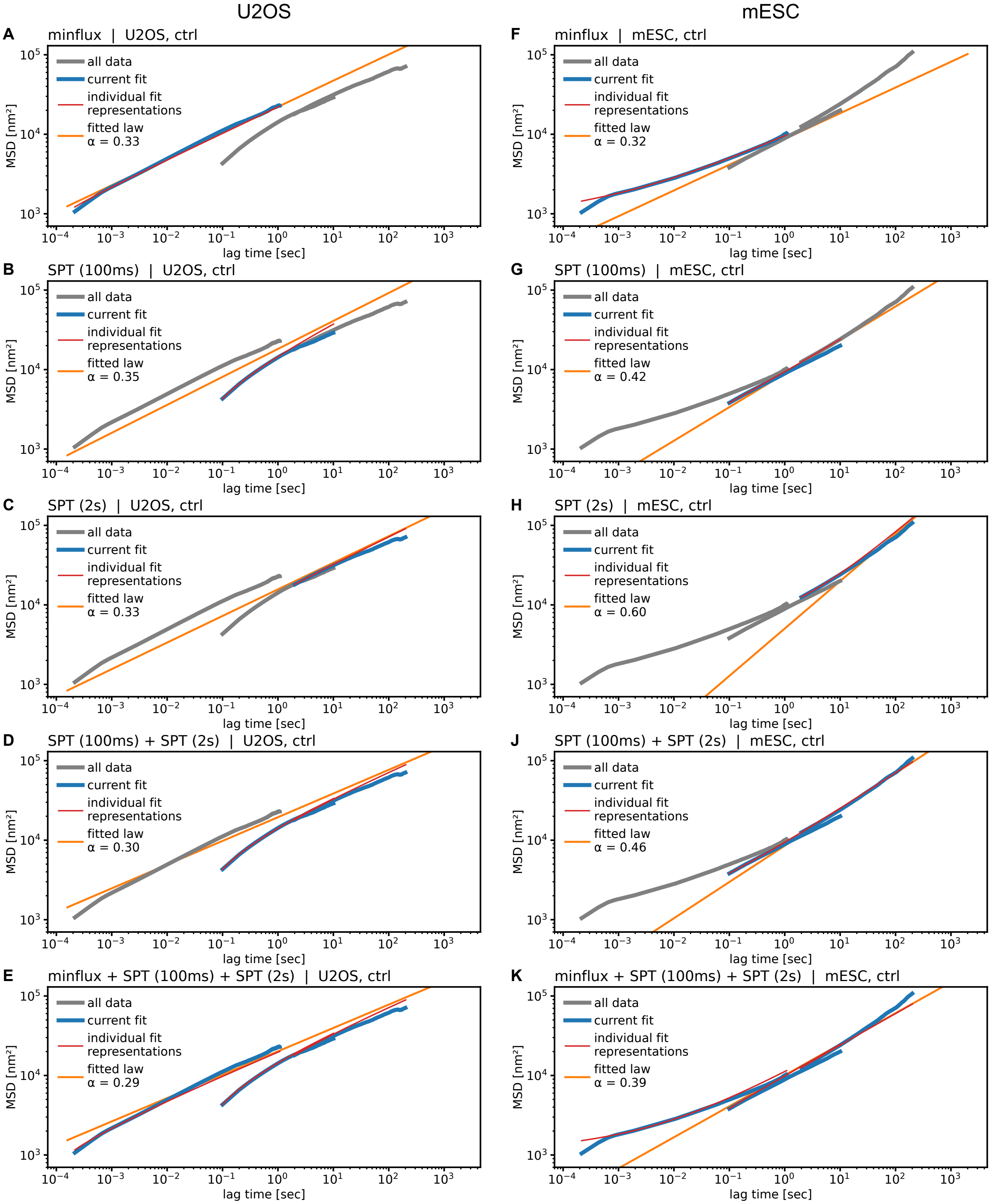

**Fig. S16:** Bayesian MSD fits to H2B tracking under control condition (DMSO) in U2OS (A–E) and mESC (F–K). Only the data set highlighted in blue (and named in the panel title) was fitted, the remaining data are shown in grey for reference. The fit result and its shape including data set specific biases (motion blur and localization error) are shown in orange and red, respectively, as described in main text Fig. 3A.

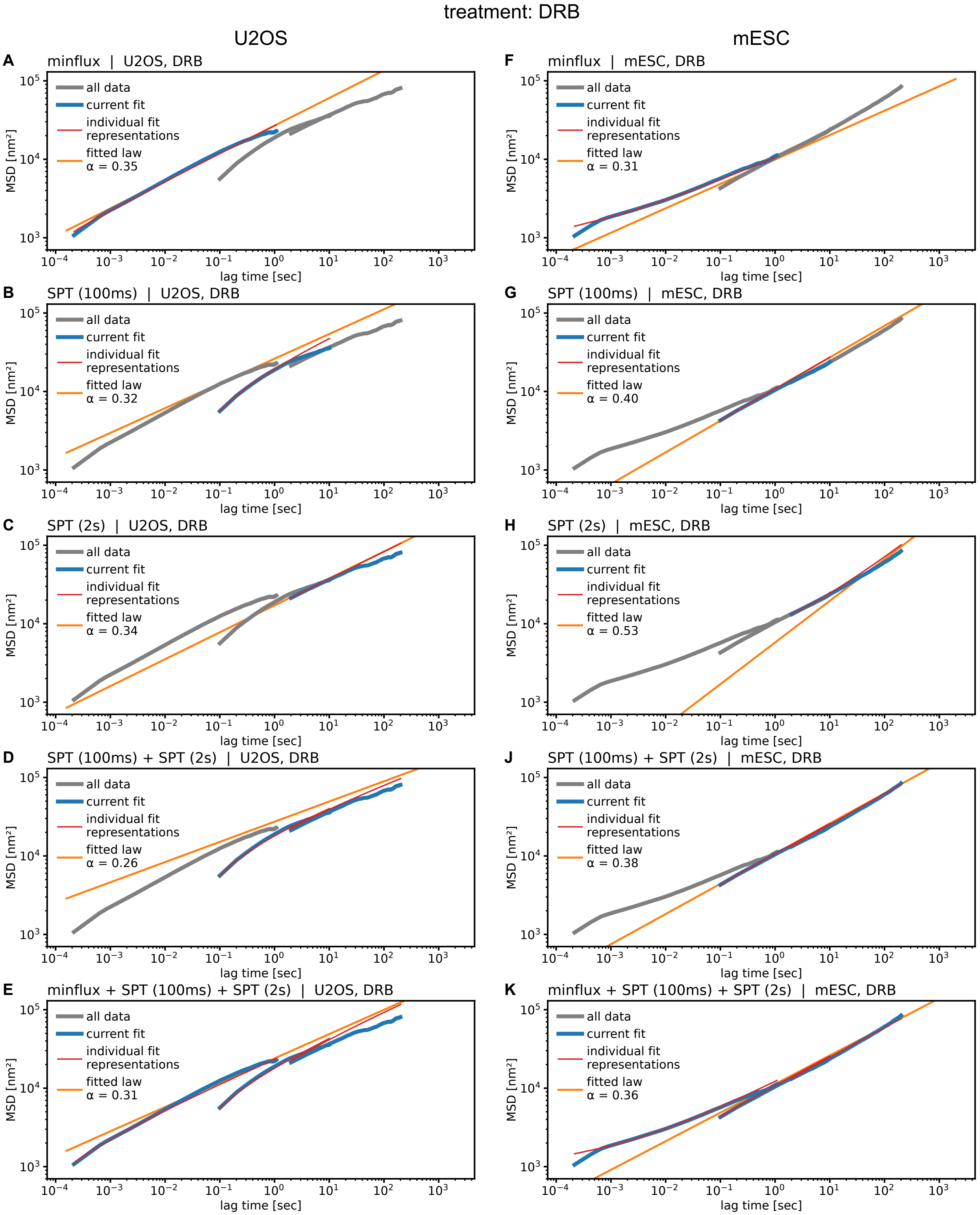

**Fig. S17:** Bayesian MSD fits to H2B tracking under DRB treatment in U2OS (A–E) and mESC (F–K). Only the data set highlighted in blue (and named in the panel title) was fitted, the remaining data are shown in grey for reference. The fit result and its shape including data set specific biases (motion blur and localization error) are shown in orange and red, respectively, as described in main text Fig. 3A.

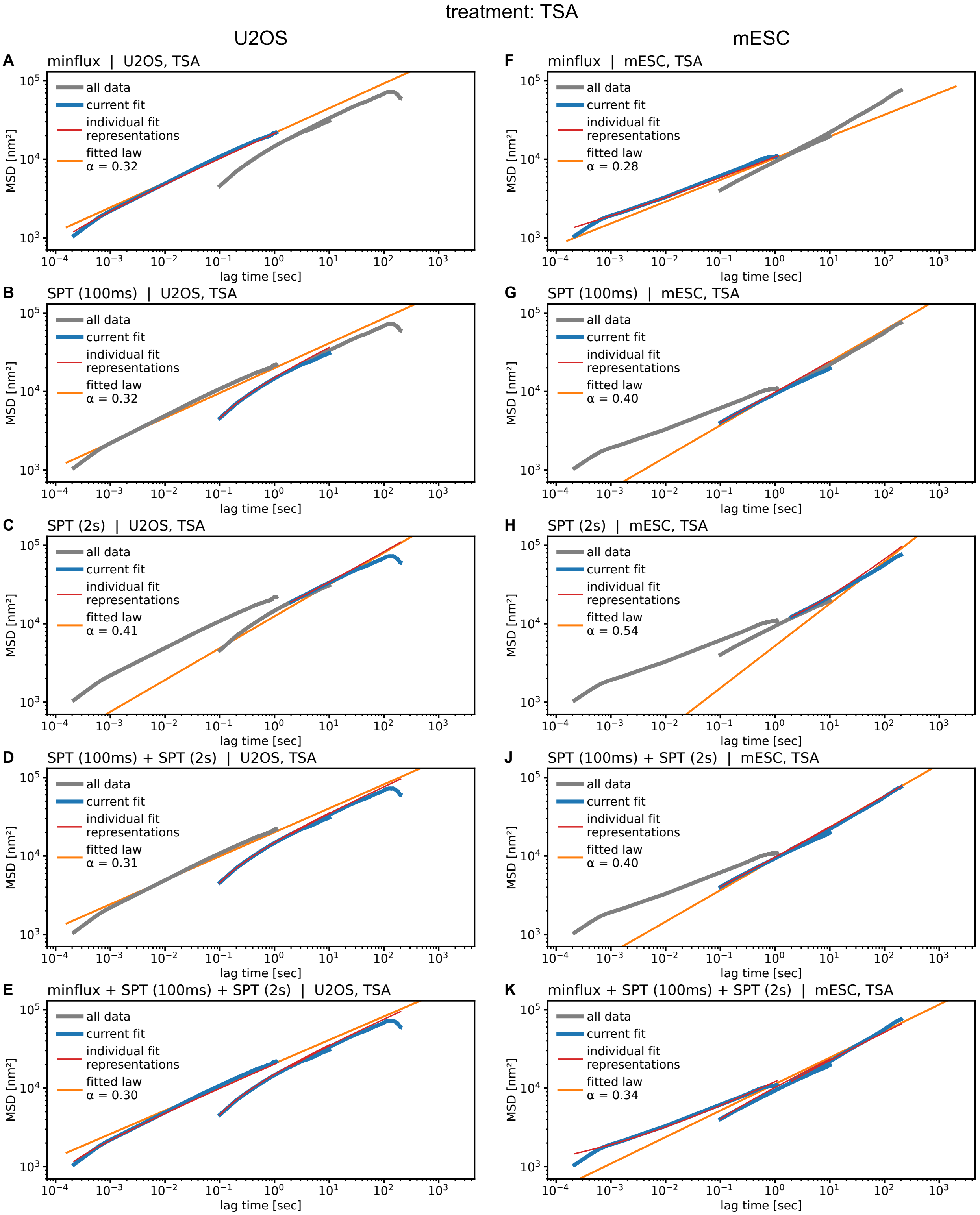

**Fig. S18:** Bayesian MSD fits to H2B tracking under TSA treatment in U2OS (A–E) and mESC (F–K). Only the data set highlighted in blue (and named in the panel title) was fitted, the remaining data are shown in grey for reference. The fit result and its shape including data set specific biases (motion blur and localization error) are shown in orange and red, respectively, as described in main text Fig. 3A.

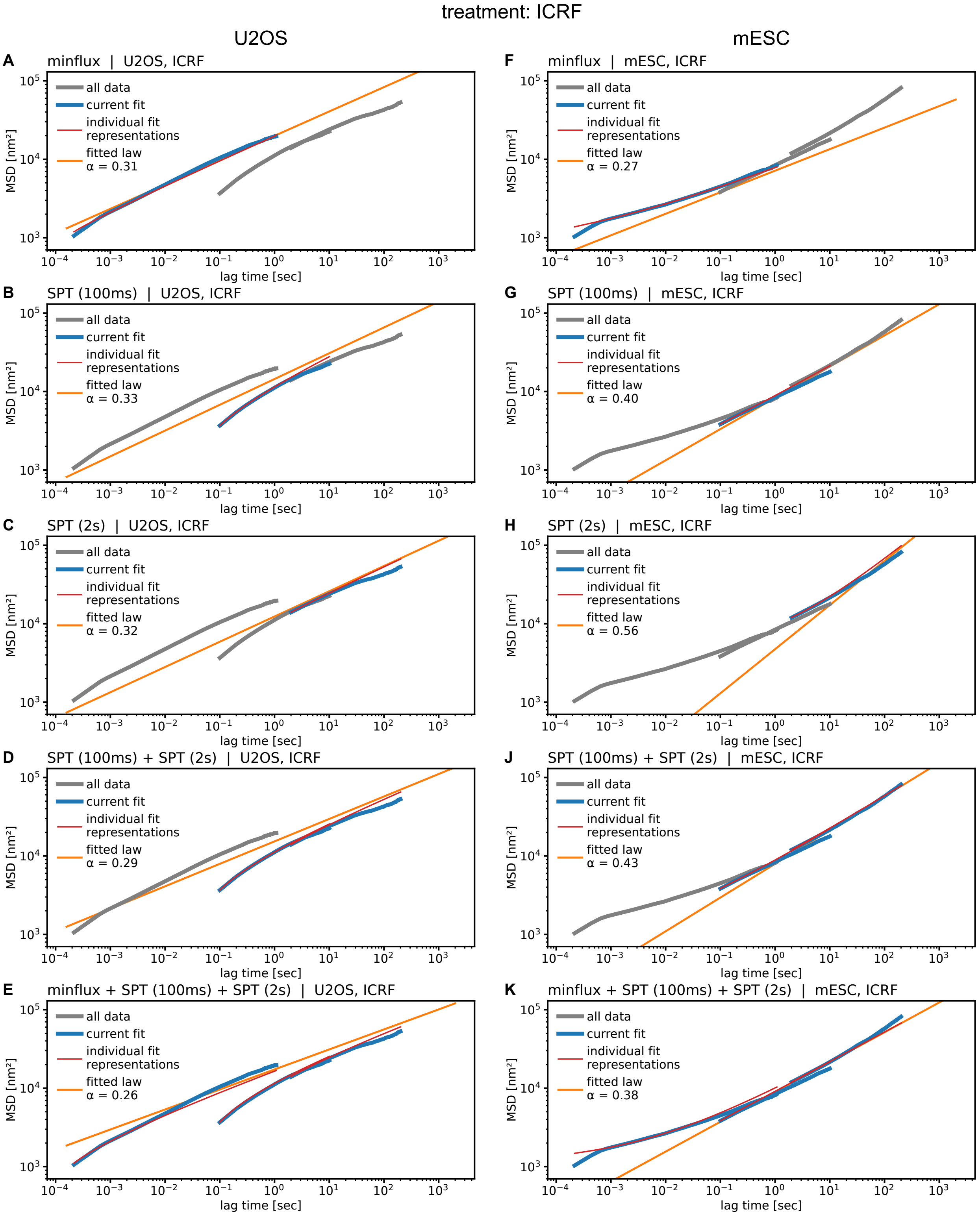

**Fig. S19:** Bayesian MSD fits to H2B tracking under ICRF treatment in U2OS (A–E) and mESC (F–K). Only the data set highlighted in blue (and named in the panel title) was fitted, the remaining data are shown in grey for reference. The fit result and its shape including data set specific biases (motion blur and localization error) are shown in orange and red, respectively, as described in main text Fig. 3A.

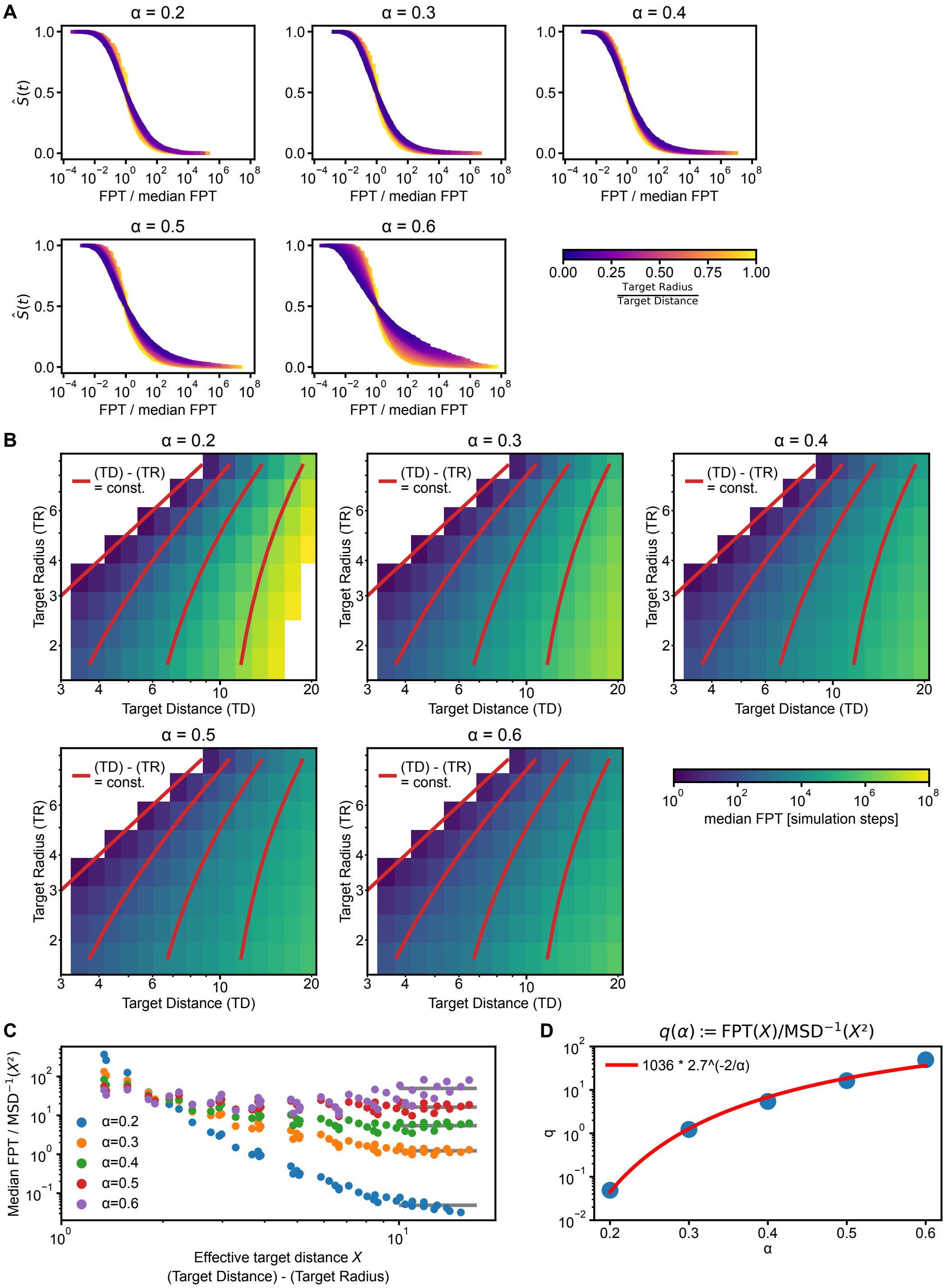

**Fig. S20:** Extracting median first passage times (FPT) from fBm simulations. (A) Kaplen-Meier survival curves for the search process, rescaled by median and color coded by relative target size. (B) Median FPT as function of target distance and radius. Missing data indicate that less than 50% of the simulated trajectories found the target, or the target volume overlapped the starting point (target radius greater than target distance). Red: level lines for "effective target distance", i.e. distance $X$ to the closest point in the target volume. (C) Median FPT rescaled by expected time scale from MSD based on effective target distance. Only data with target radius $> 3$ are shown to avoid artifacts from too small targets. Grey: data for $X > 10$ were averaged to find $q(\alpha)$. (D) Dimensionless prefactor $q$ as a function of $\alpha$. Red: a heuristic fit line.

### 7 Supplementary Tables

Table S1: MSD exponent measurements from previous studies

| **Cell type** | **Experimental system** | **MSD Time range [sec]** | **Order of magnitude**† | **No. of $\alpha$ regimes** | **Reported $\alpha$** | **Reference** |
|---|---|---|---|---|---|---|
| Mouse Embryonic Stem Cells (mESCs) | *Fbn2* locus (ANCHOR3/TetO) | $2 \times 10^1$ – $7.2 \times 10^3$ | 2 | 1 | 0.50 | (3) |
| mESCs | Multiple loci (TetO randomly integrated) | $1 \times 10^1$ – $1.8 \times 10^3$ | 2 | 1 | 0.60 | (19) |
| mESCs | *Sox2* promoter (ANCHOR1) and Control Region (ANCHOR3) | $5 \times 10^{-1}$ – $1.2 \times 10^2$ | 2 | 1 | 0.13; 0.20 | (20) |
| mESCs | Multiple loci (TetO randomly integrated) | $5 \times 10^{-1}$ – $1.2 \times 10^2$ | 2 | Distribution of $\alpha$ | 0.50[a]; 0.62[a] | (21) |
| mESCs | Multiple loci (TetO/CuO inserted at different genomic separations) | $5 \times 10^{-1}$ – $1.8 \times 10^3$ | 3 | Distribution of $\alpha$ | Variable based on genomic distance | (22) |
| mESCs | *Fgf5* locus (dCas9-eGFP) | $2 \times 10^{-1} - 8$ | 1 | 1 | 0.53 | (23) |
| Mouse epiblast-like cells | *Fgf5* locus (dCas9-eGFP) | $2 \times 10^{-1} - 8$ | 1 | 1 | 0.51 | (23) |
| Mouse B cells | VH-JH element (TetO) | $2 - 2 \times 10^3$ | 3 | 1 | 0.50 | (24) |
| Mouse B cells | DH-JH element (TetO) | $2\ 2 - 2 \times 10^3$ | 3 | 1 | 0.50 | (24) |
| Mouse B cells | VH-JH element (TetO) | $2 - 2 \times 10^3$ | 3 | 2 | 0.20; 0.35 | (25) |
| Mouse B cells | DH-JH element (mutant TetO) | $2 - 2 \times 10^3$ | 3 | 2 | 0.45; 0.70 | (25) |
| Mouse embryonic fibroblasts | Telomeres (GFP-TRF2) | $1 \times 10^{-1}$ – $1 \times 10^3$ | 4 | 2 | 0.26; 0.43 | (26) |
| NIH3T3 (mouse) | Telomeres (GFP-TRF2) | $1 \times 10^{-1}$ – $1 \times 10^3$ | 4 | 2 | 0.43; 0.47 | (26) |
| Primary mouse fibroblast | Telomeres (GFP-TRF2) | $1 \times 10^{-1}$ – $1 \times 10^3$ | 4 | 2 | 0.12; 0.76 | (26) |
| DM (Deer) | Histone H4 (H4-PA-GFP) | $3 \times 10^{-2}$ – $2 \times 10^{-1}$ | 0 | 1 | 0.31; 0.37 | (27) |

Continued on next page

Table S1: MSD exponent measurements from previous studies (Continued)

| Cell type | Experimental system | MSD Time range [sec] | Order of magnitude[†] | No. of $\alpha$ regimes | Reported $\alpha$ | Reference |
|---|---|---|---|---|---|---|
| RPE-1 (Human) | Histone H2B (H2B-Halo, H2B-PA-mCherry) | $5 \times 10^{-2} - 3$ | 1 | 1 | 0.28 | (28) |
| HT-1080 (Human) | Single locus, chr 5 (TetO) | $5 \times 10^{-2} - 3$ | 1 | 1 | 0.38 | (29) |
| HeLa (Human) | Histone H2B (H2B-PA-mCherry) | $5 \times 10^{-2}$ – $5 \times 10^{-1}$ | 1 | 4 | 0.43; 0.47; 0.68; 0.88 | (30, 31) |
| HeLa (Human) | Histone H2B (H2B-Halo) | $5 \times 10^{-2} - 5$ | 2 | 1 | 0.45 | (29) |
| HeLa (Human) | Histone H2B (H2B-Halo); DNA (Cy3-dCTP) | $5 \times 10^{-2}$ – $1 \times 10^{1}$ | 2 | 2 | 0.45; 0.52 | (32) |
| HeLa (Human) | Telomeres (GFP-TRF2) | $1 \times 10^{-1}$ – $1 \times 10^{3}$ | 4 | 2 | 0.34; 0.43 | (26) |
| U2OS (Human) | Telomeres (GFP-TRF2) | $1 \times 10^{-1}$ – $1 \times 10^{3}$ | 4 | 2 | 0.38; 0.55 | (26) |
| U2OS (Human) | Telomeres (GFP-TRF2) | $1 \times 10^{-2}$ – $1 \times 10^{4}$ | 6 | 3 | 0.32; 0.51; 1.15 | (33) |
| U2OS (Human) | Centromeres (CENPA-eGFP) | $1 \times 10^{-1}$ – $1 \times 10^{3}$ | 4 | 2 | 0.19; 0.50 | (26) |
| U2OS (Human) | Subtelomeric locus, chr 1 (TetO array) | $5 \times 10^{-1}$ – $3.6 \times 10^{3}$ | 3 | 1 | 0.50 | (34) |
| MCF7 (Human) | Single locus (ANCHOR3-CCND1-MS2) | $2.5 \times 10^{-1} - 5$ | 2 | 2 | 0.4; 0.50; 1.00 | (35) |
| Yeast | MAT locus (LacO) | $8 \times 10^{-2}$ – $2.5 \times 10^{2}$ | 3 | Distribution of $\alpha$ | 0.48[a]; 0.62[a] | (36) |
| Yeast | Single locus, chr XII (TetO) | $2 \times 10^{-1}$ – $1.2 \times 10^{1}$ | 1 | 1 | 0.54 | (37) |
| Yeast | Multiple loci, chr III, IV, XII, and XIV (TetO) | $2 \times 10^{-1}$ – $1.4 \times 10^{2}$ | 2 | 1 | 0.52 | (38) |
| Drosophila | *eve* locus (MS2) and distal target gene (parS) | $3 \times 10^{1} - 1 \times 10^{3}$ | 2 | 1 | 0.24 | (39) |
| C. elegans | Single locus (LacO) | $2 \times 10^{1}$ – $1.5 \times 10^{2}$ | 1 | Distribution of $\alpha$ | 0.30 – 0.50[b] | (40) |

[†] Values are rounded down to the nearest whole number.
[a] Average value reported in the paper.
[b] Lower and upper values of the distribution range.

Table S2: Bayesian fit results

| Cell type | Treatment | Data set | Fit parameter | value | 95% credible interval |
|---|---|---|---|---|---|
| U2OS | ctrl | MINFLUX | $\alpha$ | 0.329 | [ 0.324, 0.334] |
| | | | $\log(\alpha\Gamma)$ | -5.614 | [ -5.643, -5.585] |
| | | | $\sigma$ MINFLUX | 13.8 nm | [ 13.6, 14.0] nm |
| | | SPT 100ms | $\alpha$ | 0.352 | [ 0.340, 0.364] |
| | | | $\log(\alpha\Gamma)$ | -5.744 | [ -5.766, -5.722] |
| | | | $\sigma$ SPT-100ms | 17.3 nm | [ 16.8, 17.8] nm |
| | | SPT 2s | $\alpha$ | 0.335 | [ 0.313, 0.356] |
| | | | $\log(\alpha\Gamma)$ | -5.943 | [ -5.984, -5.903] |
| | | | $\sigma$ SPT-2s | 26.3 nm | [ 21.0, 30.4] nm |
| | | SPT (both) | $\alpha$ | 0.298 | [ 0.293, 0.304] |
| | | | $\log(\alpha\Gamma)$ | -5.843 | [ -5.850, -5.835] |
| | | | $\sigma$ SPT-100ms | 15.6 nm | [ 15.2, 15.9] nm |
| | | | $\sigma$ SPT-2s | 10.6 nm | [ 8.1, 12.6] nm |
| | | MINFLUX + SPT | $\alpha$ | 0.294 | [ 0.293, 0.295] |
| | | | $\log(\alpha\Gamma)$ | -5.823 | [ -5.829, -5.816] |
| | | | $\sigma$ SPT-100ms | 14.7 nm | [ 14.5, 15.0] nm |
| | | | $\sigma$ SPT-2s | 0.7 nm | [ 0.0, 6.7] nm |
| | | | $\sigma$ MINFLUX | 12.7 nm | [ 12.6, 12.8] nm |
| | DRB | MINFLUX | $\alpha$ | 0.352 | [ 0.347, 0.357] |
| | | | $\log(\alpha\Gamma)$ | -5.354 | [ -5.385, -5.322] |
| | | | $\sigma$ MINFLUX | 13.4 nm | [ 13.2, 13.6] nm |
| | | SPT 100ms | $\alpha$ | 0.315 | [ 0.303, 0.327] |
| | | | $\log(\alpha\Gamma)$ | -5.491 | [ -5.513, -5.468] |
| | | | $\sigma$ SPT-100ms | 16.7 nm | [ 15.9, 17.4] nm |
| | | SPT 2s | $\alpha$ | 0.343 | [ 0.315, 0.371] |
| | | | $\log(\alpha\Gamma)$ | -5.829 | [ -5.880, -5.777] |
| | | | $\sigma$ SPT-2s | 32.9 nm | [ 26.8, 37.5] nm |
| | | SPT (both) | $\alpha$ | 0.257 | [ 0.252, 0.263] |
| | | | $\log(\alpha\Gamma)$ | -5.650 | [ -5.659, -5.641] |
| | | | $\sigma$ SPT-100ms | 15.5 nm | [ 15.0, 16.1] nm |
| | | | $\sigma$ SPT-2s | 0.0 nm | [ 0.0, 2.2] nm |
| | | MINFLUX + SPT | $\alpha$ | 0.310 | [ 0.308, 0.311] |
| | | | $\log(\alpha\Gamma)$ | -5.596 | [ -5.603, -5.588] |
| | | | $\sigma$ SPT-100ms | 19.0 nm | [ 18.7, 19.2] nm |
| | | | $\sigma$ SPT-2s | 0.0 nm | [ 0.0, 2.3] nm |
| | | | $\sigma$ MINFLUX | 11.8 nm | [ 11.6, 11.9] nm |
| | TSA | MINFLUX | $\alpha$ | 0.316 | [ 0.311, 0.321] |
| | | | $\log(\alpha\Gamma)$ | -5.678 | [ -5.708, -5.649] |
| | | | $\sigma$ MINFLUX | 13.3 nm | [ 13.2, 13.5] nm |

Continued on next page

Table S2: Bayesian fit results (Continued)

| Cell type | Treatment | Data set | Fit parameter | value | 95% credible interval |
|---|---|---|---|---|---|
| | | SPT 100ms | $\alpha$ | 0.317 | [ 0.303, 0.331] |
| | | | $\log(\alpha\Gamma)$ | -5.759 | [ -5.784, -5.733] |
| | | | $\sigma$ SPT-100ms | 16.9 nm | [ 16.3, 17.6] nm |
| | | SPT 2s | $\alpha$ | 0.406 | [ 0.384, 0.426] |
| | | | $\log(\alpha\Gamma)$ | -5.983 | [ -6.016, -5.947] |
| | | | $\sigma$ SPT-2s | 36.2 nm | [ 33.7, 38.3] nm |
| | | SPT (both) | $\alpha$ | 0.306 | [ 0.300, 0.311] |
| | | | $\log(\alpha\Gamma)$ | -5.789 | [ -5.797, -5.783] |
| | | | $\sigma$ SPT-100ms | 16.7 nm | [ 16.4, 17.1] nm |
| | | | $\sigma$ SPT-2s | 11.1 nm | [ 8.5, 13.2] nm |
| | | MINFLUX + SPT | $\alpha$ | 0.300 | [ 0.298, 0.301] |
| | | | $\log(\alpha\Gamma)$ | -5.777 | [ -5.783, -5.771] |
| | | | $\sigma$ SPT-100ms | 16.0 nm | [ 15.7, 16.2] nm |
| | | | $\sigma$ SPT-2s | 5.8 nm | [ 0.0, 9.0] nm |
| | | | $\sigma$ MINFLUX | 12.8 nm | [ 12.7, 13.0] nm |
| | ICRF | MINFLUX | $\alpha$ | 0.310 | [ 0.305, 0.316] |
| | | | $\log(\alpha\Gamma)$ | -5.779 | [ -5.811, -5.748] |
| | | | $\sigma$ MINFLUX | 13.5 nm | [ 13.3, 13.7] nm |
| | | SPT 100ms | $\alpha$ | 0.329 | [ 0.319, 0.339] |
| | | | $\log(\alpha\Gamma)$ | -6.044 | [ -6.062, -6.026] |
| | | | $\sigma$ SPT-100ms | 17.3 nm | [ 16.9, 17.6] nm |
| | | SPT 2s | $\alpha$ | 0.322 | [ 0.303, 0.341] |
| | | | $\log(\alpha\Gamma)$ | -6.219 | [ -6.252, -6.184] |
| | | | $\sigma$ SPT-2s | 23.6 nm | [ 19.6, 26.6] nm |
| | | SPT (both) | $\alpha$ | 0.286 | [ 0.282, 0.291] |
| | | | $\log(\alpha\Gamma)$ | -6.121 | [ -6.128, -6.114] |
| | | | $\sigma$ SPT-100ms | 16.2 nm | [ 15.9, 16.4] nm |
| | | | $\sigma$ SPT-2s | 10.9 nm | [ 9.2, 12.3] nm |
| | | MINFLUX + SPT | $\alpha$ | 0.255 | [ 0.254, 0.257] |
| | | | $\log(\alpha\Gamma)$ | -6.115 | [ -6.120, -6.109] |
| | | | $\sigma$ SPT-100ms | 13.9 nm | [ 13.7, 14.1] nm |
| | | | $\sigma$ SPT-2s | 0.1 nm | [ 0.0, 3.0] nm |
| | | | $\sigma$ MINFLUX | 11.8 nm | [ 11.7, 12.0] nm |
| mESC | ctrl | MINFLUX | $\alpha$ | 0.324 | [ 0.316, 0.331] |
| | | | $\log(\alpha\Gamma)$ | -6.563 | [ -6.603, -6.522] |
| | | | $\sigma$ MINFLUX | 17.7 nm | [ 17.6, 17.8] nm |
| | | SPT 100ms | $\alpha$ | 0.422 | [ 0.411, 0.434] |
| | | | $\log(\alpha\Gamma)$ | -6.277 | [ -6.295, -6.258] |

Continued on next page

Table S2: Bayesian fit results (Continued)

| Cell type | Treatment | Data set | Fit parameter | value | 95% credible interval |
|---|---|---|---|---|---|
| | | | $\sigma$ SPT-100ms | 24.3 nm | [ 24.1, 24.5] nm |
| | | SPT 2s | $\alpha$ | 0.596 | [ 0.579, 0.613] |
| | | | $\log(\alpha\Gamma)$ | -6.501 | [ -6.528, -6.469] |
| | | | $\sigma$ SPT-2s | 37.2 nm | [ 36.1, 38.1] nm |
| | | SPT (both) | $\alpha$ | 0.456 | [ 0.451, 0.461] |
| | | | $\log(\alpha\Gamma)$ | -6.244 | [ -6.252, -6.236] |
| | | | $\sigma$ SPT-100ms | 24.9 nm | [ 24.7, 25.0] nm |
| | | | $\sigma$ SPT-2s | 24.7 nm | [ 24.1, 25.3] nm |
| | | MINFLUX + SPT | $\alpha$ | 0.390 | [ 0.388, 0.392] |
| | | | $\log(\alpha\Gamma)$ | -6.232 | [ -6.238, -6.224] |
| | | | $\sigma$ SPT-100ms | 23.1 nm | [ 23.0, 23.2] nm |
| | | | $\sigma$ SPT-2s | 20.5 nm | [ 19.8, 21.2] nm |
| | | | $\sigma$ MINFLUX | 18.5 nm | [ 18.5, 18.6] nm |
| | DRB | MINFLUX | $\alpha$ | 0.311 | [ 0.304, 0.319] |
| | | | $\log(\alpha\Gamma)$ | -6.472 | [ -6.515, -6.430] |
| | | | $\sigma$ MINFLUX | 17.1 nm | [ 17.0, 17.2] nm |
| | | SPT 100ms | $\alpha$ | 0.403 | [ 0.392, 0.414] |
| | | | $\log(\alpha\Gamma)$ | -6.135 | [ -6.153, -6.117] |
| | | | $\sigma$ SPT-100ms | 25.0 nm | [ 24.8, 25.2] nm |
| | | SPT 2s | $\alpha$ | 0.531 | [ 0.515, 0.547] |
| | | | $\log(\alpha\Gamma)$ | -6.482 | [ -6.512, -6.460] |
| | | | $\sigma$ SPT-2s | 37.8 nm | [ 36.7, 38.8] nm |
| | | SPT (both) | $\alpha$ | 0.385 | [ 0.380, 0.389] |
| | | | $\log(\alpha\Gamma)$ | -6.186 | [ -6.191, -6.178] |
| | | | $\sigma$ SPT-100ms | 25.0 nm | [ 24.8, 25.1] nm |
| | | | $\sigma$ SPT-2s | 21.0 nm | [ 20.3, 21.7] nm |
| | | MINFLUX + SPT | $\alpha$ | 0.364 | [ 0.363, 0.366] |
| | | | $\log(\alpha\Gamma)$ | -6.187 | [ -6.192, -6.181] |
| | | | $\sigma$ SPT-100ms | 24.4 nm | [ 24.2, 24.5] nm |
| | | | $\sigma$ SPT-2s | 19.7 nm | [ 19.0, 20.4] nm |
| | | | $\sigma$ MINFLUX | 17.8 nm | [ 17.8, 17.9] nm |
| | TSA | MINFLUX | $\alpha$ | 0.277 | [ 0.269, 0.285] |
| | | | $\log(\alpha\Gamma)$ | -6.549 | [ -6.592, -6.506] |
| | | | $\sigma$ MINFLUX | 16.3 nm | [ 16.2, 16.5] nm |
| | | SPT 100ms | $\alpha$ | 0.405 | [ 0.395, 0.415] |
| | | | $\log(\alpha\Gamma)$ | -6.262 | [ -6.278, -6.246] |
| | | | $\sigma$ SPT-100ms | 24.9 nm | [ 24.7, 25.0] nm |
| | | SPT 2s | $\alpha$ | 0.538 | [ 0.525, 0.550] |
| | | | $\log(\alpha\Gamma)$ | -6.568 | [ -6.590, -6.549] |

Continued on next page

Table S2: Bayesian fit results (Continued)

| **Cell type** | **Treatment** | **Data set** | **Fit parameter** | **value** | **95% credible interval** |
|---|---|---|---|---|---|
| | | | $\sigma$ SPT-2s | 37.0 nm | [ 36.4, 37.6] nm |
| | | SPT (both) | $\alpha$ | 0.401 | [ 0.397, 0.405] |
| | | | $\log(\alpha\Gamma)$ | -6.295 | [ -6.301, -6.288] |
| | | | $\sigma$ SPT-100ms | 25.0 nm | [ 24.9, 25.1] nm |
| | | | $\sigma$ SPT-2s | 23.4 nm | [ 22.9, 23.9] nm |
| | | MINFLUX + SPT | $\alpha$ | 0.338 | [ 0.336, 0.340] |
| | | | $\log(\alpha\Gamma)$ | -6.263 | [ -6.267, -6.258] |
| | | | $\sigma$ SPT-100ms | 22.9 nm | [ 22.8, 23.0] nm |
| | | | $\sigma$ SPT-2s | 17.3 nm | [ 16.7, 17.9] nm |
| | | | $\sigma$ MINFLUX | 17.6 nm | [ 17.5, 17.7] nm |
| | ICRF | MINFLUX | $\alpha$ | 0.275 | [ 0.267, 0.282] |
| | | | $\log(\alpha\Gamma)$ | -6.925 | [ -6.967, -6.883] |
| | | | $\sigma$ MINFLUX | 17.1 nm | [ 17.0, 17.2] nm |
| | | SPT 100ms | $\alpha$ | 0.398 | [ 0.387, 0.409] |
| | | | $\log(\alpha\Gamma)$ | -6.406 | [ -6.423, -6.389] |
| | | | $\sigma$ SPT-100ms | 24.8 nm | [ 24.6, 25.0] nm |
| | | SPT 2s | $\alpha$ | 0.562 | [ 0.546, 0.579] |
| | | | $\log(\alpha\Gamma)$ | -6.620 | [ -6.651, -6.590] |
| | | | $\sigma$ SPT-2s | 37.5 nm | [ 36.5, 38.5] nm |
| | | SPT (both) | $\alpha$ | 0.430 | [ 0.425, 0.435] |
| | | | $\log(\alpha\Gamma)$ | -6.374 | [ -6.382, -6.366] |
| | | | $\sigma$ SPT-100ms | 25.3 nm | [ 25.2, 25.4] nm |
| | | | $\sigma$ SPT-2s | 27.0 nm | [ 26.5, 27.5] nm |
| | | MINFLUX + SPT | $\alpha$ | 0.380 | [ 0.378, 0.382] |
| | | | $\log(\alpha\Gamma)$ | -6.376 | [ -6.383, -6.368] |
| | | | $\sigma$ SPT-100ms | 24.1 nm | [ 24.0, 24.2] nm |
| | | | $\sigma$ SPT-2s | 25.0 nm | [ 24.5, 25.5] nm |
| | | | $\sigma$ MINFLUX | 18.3 nm | [ 18.3, 18.4] nm |

#### 8 Supplementary Movies

Representative 2-color SPT movies for H2B-Halo, acquired in mESCs or U2OS cells. As described in section 2.1, prior to SPT acquisition, we performed 2-color labeling using different concentrations of the Halo dyes $JFX_{554}$ to obtain reference images of nuclei for segmentation, and $JFX_{650}$ to track H2B-Halo by SPT.

Each movie therefore includes: (1) A reference image of the nucleus, obtained by exciting H2B-Halo with a 561 nm laser for one frame (Frame 1 in the raw movies). For representation purposes, this frame is shown here for 20 consecutive frames. (2) SPT data acquired by exciting H2B-Halo with a 640 nm laser over 450 frames. (3) A final reference image of the nucleus, acquired at the end of the movie (Frame 452 in the raw movies). Like the initial reference, it is shown for 20 consecutive frames.

All movies are displayed with overlaid ROIs, manually drawn prior to SPT tracking (see section 2.2), and shown with adjusted contrast and rolling ball background subtraction (*41*) (radius: 20 pixels) for representation purposes only.

**Supplementary Movie 1.** Representative SPT movie for H2B-Halo, in mESCs, collected at 100 ms frame interval (10 fps) and displayed at 10 fps.

**Supplementary Movie 2.** Representative SPT movie for H2B-Halo, in U2OS, collected at 100 ms frame interval (10 fps) and displayed at 10 fps.

**Supplementary Movie 3.** Representative SPT movie for H2B-Halo, in mESCs, collected at 2 sec frame interval (0.5 fps) and displayed at 10 fps.

**Supplementary Movie 4.** Representative SPT movie for H2B-Halo, in U2OS, collected at 2 sec frame interval (0.5 fps) and displayed at 10 fps.
